# Eco-genomic analysis uncovers precision-conservation targets for the western Pacific’s southernmost salmonid

**DOI:** 10.1101/2025.09.09.675276

**Authors:** Yi-Chien Lee, Zong-Yu Shen, Wei-Ren Lin, Tzi-Yuan Wang, Mark Liu, Yu-Chen Yeh, Shih-Fan Chan, Mei-Yeh Jade Lu, Hui-Yu Wang, Lin-Yan Liao, Wen-Hsiung Li, Jen-Pan Huang, Isheng Jason Tsai, Sheng-Feng Shen

## Abstract

Understanding how isolated small populations persist and adapt in adverse environments is instrumental to evolutionary and conservation biology. We combine a chromosome-level genome assembly, population resequencing and forward-time simulations to reconstruct the history and viability of the Formosan landlocked salmon (*Oncorhynchus formosanus*), now restricted to a handful of high-mountain headwaters in Taiwan. We estimate that this lineage has diverged from Japanese masu salmon over one million years ago and has no detectable gene flow for ∼50,000 years. It has accumulated extensive chromosome fusions and expansions of cold-adaptation gene families, qualifying it as a new species rather than a subspecies of Japanese masu salmon. Whole-stream sampling reveals an overlooked Hehuan-Creek population that retains high heterozygosity and has gained unique alleles. Life-table simulations show that the Hehuan population has a notably lower extinction risk and can persist or even grow under low-to-moderate typhoon frequency, whereas Qijiawan-Creek population would decline precipitously under the same or higher frequency . These findings contradict the notion that peripheral populations are likely genetically depleted and support stream-specific “precision conservation” in place of broad, untargeted translocations that could erode local adaptation potential. Thus, our genomic-ecological analysis has uncovered hidden strong resilience in a critically endangered, climate-threatened salmonid lineage.

## Introduction

How do small populations at the margins of a species’ range maintain genetic diversity and adaptive potential under long-term isolation and extreme environmental conditions^1^? Moreover, what are the implications of these dynamics in the face of accelerating climate change^2,3^? These questions lie at the heart of modern conservation biology. Marginal populations, though often small and fragmented, may harbor unique adaptive variation shaped by intense local selection^4^. For example, edge populations of Australian vertebrates in hot, arid regions have evolved enhanced drought or heat tolerance, potentially serving as reservoirs of climate-relevant alleles^5^. In salmonids, local adaptation is both rapid and fine-scaled: within just a few generations, native populations frequently outperform immigrants in survival and reproductive success^6^.

However, evolutionary outcomes are not dictated by selection alone. In very small populations, genetic drift can overwhelm selection, eroding adaptive variation and fixing deleterious alleles^7,8^. Migrants may occasionally restore lost variation or even outperform residents^9^. This interplay between drift and selection complicates conservation decisions. Peripheral populations are often recognized as evolutionarily significant units (ESUs) for their unique genetic legacies^10^, but also face elevated extinction risks due to inbreeding, reduced heterozygosity, and loss of adaptive potential^3,11,12^. Genetic rescue—introducing individuals from other populations to restore diversity—is increasingly promoted^9,13^, though not without risks of outbreeding depression^14^ and sociopolitical resistance, especially in culturally iconic taxa.

The Formosan landlocked salmon (*Oncorhynchus formosanus*, Jordan & Oshima 1919) exemplifies this conservation paradox. Endemic to Taiwan, it is the southernmost lineage and one of the most tropical salmonids, classified within the *O. masou* species complex alongside the Japanese Masu (*O. m. masou*), Amago (*O. m. ishikawae*)^15^, and Biwa salmon—now recognized as the distinct species *O. biwaensis*^16^ rather than *O. m. rhodurus* (Fig 1a). Morphologically distinct and geographically isolated by over 1,000 km of ocean and mountains^17^, its taxonomic status remains contested. Although the IUCN lists it as a distinct and Critically Endangered species^18^, genetic studies have often placed it within the *O. masou* clade, based on limited molecular data^15,19–21^. Although species-level recognition has been proposed^22^, this view was based on earlier datasets, lacking new genetic evidence.

**Figure 1.**
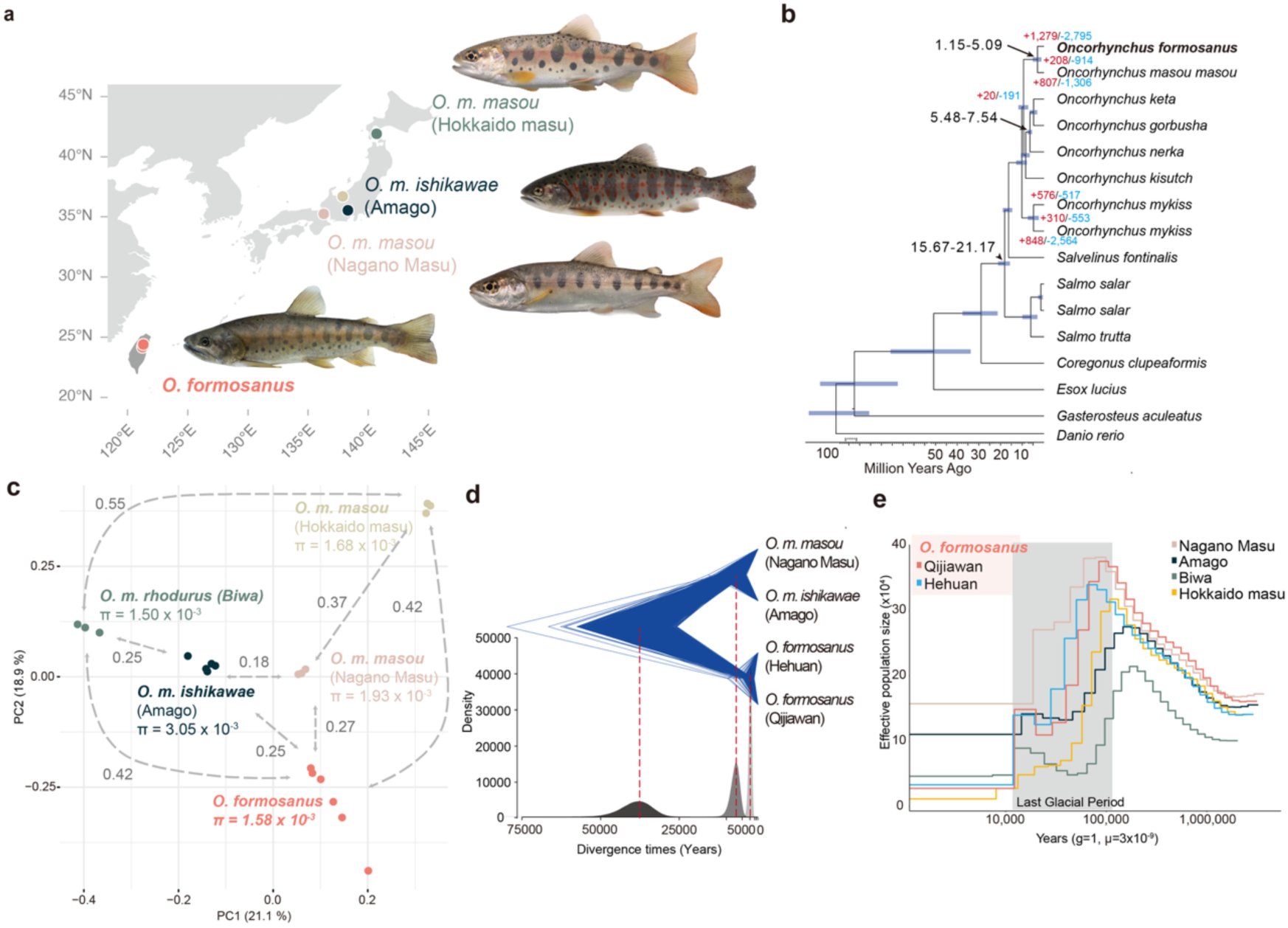
Phylogenomics and demographic history of *O. formosanus*. (**a**) Geographic distribution and phenotypic variation of selected lineages within the *Oncorhynchus masou* species complex. (**b**) Species tree inferred from a concatenated alignment of 1,483 single-copy orthogroups. Blue horizontal bars representing the 95% confidence intervals. Numbers in red and blue denote the gained and lost orthogroups at each node, respectively. (**c**) Principal component analysis (PCA) showing genetic differentiation among masu salmon lineages. Numbers along dashed arrows represent mean pairwise homozygous SNP differences (in millions) between populations. (d) Time-calibrated diversification within Formosan (*O. formosanus*) and Japanese (Nagano Masu and Amago) salmon lineages visualized by DensiTree, indicating lineage splits occurring approximately 50,000–20,000 years ago. (e) Demographic histories inferred by PSMC, showing changes in effective population size for Japanese and Taiwanese salmon individuals.

This taxonomic ambiguity complicates conservation efforts. After decades of habitat degradation and overharvesting, wild populations fell to ∼600 individuals by the 1980s^23^. Conservation actions, including captive breeding and habitat restoration, gradually increased the population to ∼1,000 individuals. However, at that time the species remained confined to a single confirmed stronghold—a 7-km stretch of Chichiawan (Qijiawan) Creek, with the Luoyewei population being hatchery-supplemented and the Hehuan population presumed extirpated since the 1960s (Fig. S1). In 2017, reintroduction began in Hehuan Creek using juveniles bred from Qijiawan-derived broodstock. Over three years, more than 6,000 juveniles were released across three sectors of Hehuan Creek^24,25^. However, post-reintroduction assessments have focused narrowly on census numbers, with limited attention to genetic monitoring. Early molecular studies identified low diversity, likely reflecting historical bottlenecks and long-term isolation, but these inferences relied on sparse sampling and low-resolution markers. The extent of genome-wide variation and its relevance to adaptation remains poorly characterized. More recently, a controversial proposal suggested hybridizing *O. formosanus* with Japanese masu salmon to boost genetic diversity and resilience^26^. This proposal lacks comprehensive genomic validation and remains divisive, particularly given the species’ iconic status in Taiwan. Much of the controversy stems from insufficient genomic data, hampering rigorous assessment of the risks and benefits of such genetic rescue interventions.

In this study, we integrate whole-genome sequencing, extensive population sampling, demographic modeling, and ecological monitoring to reevaluate the evolutionary legacy and conservation outlook of *O. formosanus*, a critically endangered salmonid persisting at the tropical edge of its range. We assess whether this long-isolated lineage retains meaningful adaptive potential despite historical bottlenecks, and how genomic and demographic patterns vary across relictual stream populations. By linking genome-wide diversity with stream-specific life-history dynamics and climate-linked disturbances, we uncover mechanisms of persistence—and vulnerability—that have broader relevance for managing small, fragmented populations in a rapidly changing world.

## Results

### Sequencing, assembly and annotation of the *O. formosanus* genome

We sequenced the genome of *O. formosanus* from a single female individual from the Qijiawan Creek, one of the last remaining habitats. An initial assembly of ∼2.9 Gb was generated from 50.0x PacBio HiFi reads (Table S1), similar to genome sizes reported for other salmonids (Table S2). This assembly was further scaffolded with 31.4x Omni-C reads using YaHS^27^ (Fig. S2). The final chromosome-level assembly anchored 97.3% of the genome onto 25 scaffolds (Table S2), representing another high-quality genomic resource within the *Oncorhynchus* species; the assembly represents one of the fewest salmon chromosome numbers^28^. In addition, the mitochondrial genome was separately assembled into a single circular contig of 16.6kb^29^. The *O. formosanus* genome exhibits 56-65% repetitive content per chromosome, predominantly comprising the DNA transposon family (Table S3), like other salmonids^30,31^.

Using a combination of transcriptome sequencing from multiple tissues and proteomes from eight representative salmonids as references, we predicted 43,110 coding genes (Methods) in *O. formosanus*, which was estimated to be 95.7% complete based on BUSCO (benchmarking universal single-copy orthologs)^32^ assessment. Of these predictions, 84.1% could be assigned to 6,696 protein families using Pfam scan and 89.3% could be assigned Gene Ontology terms using eggNOG-mapper^33^. Orthology analysis with Orthofinder^34^ placed these predicted gene models alongside protein sequences from eleven representative salmonid and four outgroup genomes (Table S4) into 30,011 orthologous groups (OGs),97.7% of which were shared broadly across salmonid species, and only 0.45% were specific to *O. formosanus*. We mapped genomic sequencing data from three to five representative individuals of Japanese Amago, Biwa, Nagano and Hakkaido Masu, as well as six individuals from different Taiwanese populations, to assess genomic diversity within and between lineages (Table S1). Sequencing depth ranged from 18 - 41.8x coverage, with high mapping rates (99.49% to 99.74%;) achieved for all samples against our *O. formosanus* assembly compared to the existing *O. m. masou* reference genome, indicating close genomic affinity and reliable alignment quality (Table S5). Further genomic diversity was assessed using double-digest restriction-site-associated DNA sequencing (ddRADseq) data generated from 21, 17, and 40 individuals collected from Qijiawan, Luoyewei, and Hehuan creek populations, respectively (Table S1).

### Delimitation and demography of the *Oncorhynchus masou* species complex

We conducted phylogenomic analyses using 1,484 genome-wide single-copy ortholog sets to infer the evolutionary history and taxonomic position of *O. formosanus* relative to other salmonids. Maximum likelihood analyses of a concatenated supermatrix and coalescent-based species tree analyses placed *O. formosanus* within the *Oncorhynchus masou* species complex as the sister lineage to *O. m. masou* (Fig. 1a and Fig. S3). Using MCMCtree^35^ with two fossil calibrations^36^, we estimated the most recent common ancestor of these two lineages to be 1.2 to 5.1 million years ago (Mya) during the Pliocene and Pleistocene epoch (Fig. 1b), indicating long term evolutionary isolation. Principal component analysis (PCA) based on genome-wide SNP data clearly separated different taxa from the previously designated masu salmon species complex along two primary axes (Fig. 1c). The Biwa (*O. m. rhodurus*) lineage was distinctly differentiated along PC1, while the divergence between *O. formosanus* and *O. m. masou* appeared prominently along PC2, consistent with their geographic extremes (Central Taiwan versus Hokkaido, Japan). Pairwise genetic differentiation Fst was highest among the geographically isolated lineages (Biwa, *O. m. masou* and *O. formosanus*, Fig. S4), whereas lower differentiation among geographically southern lineages (Amago (*O. m. ishikawae)*, Nagano Masu, and *O. formosanus*; 0.18–0.27) suggests relatively recent divergence or historical gene flow among these lineages (Fig. S4).

To statistically delimit the taxonomic status of the Taiwanese *O. formosanus* and two Japanese southern lineages within the *O. masou* species complex, we first reconstructed a species tree among the three taxa based on a multi-species coalescent model^37^ and the ddRADseq data sets (Fig. 1d): the earliest divergence (∼50,000 years ago) separated the Taiwanese lineages (Hehuan and Qijiawan) from the Japanese lineages (Nagano Masu and Amago), followed by the divergence between Nagano Masu and Amago ∼ 20,000 years ago. By using a species delimitation model while allowing the species tree to vary among MCMC searches (A11 model; Methods), our genome-wide data sets suggested three species, or three evolutionarily independent lineages^38^. Specifically, the two populations within Taiwan belong to the same evolutionary lineage, which is independent from the other two evolutionary lineages, Nagano Masu and Amago. The support for three independent lineages in our data sets received > 99% posterior probability. In addition, the genealogical divergence index (GDI) estimated from our genome-wide data sets further supported substantial differentiation between Taiwanese and Japanese salmon lineages, with notably high values (Fig. S5), exceeding previously reported thresholds distinguishing biological species^39,40^. The topology of the maximum likelihood phylogeny from both the nuclear and mitochondrial genome trees also placed the *O. formosanus* species sister to all Japanese lineages (Fig. S6), suggesting an ancient split of Taiwanese salmons well before the diversification of regional populations in the Japanese archipelago. Together, these results strongly indicate that *O. formosanus* represents a distinct evolutionary significant unit (ESU) separate from the Japanese masu salmon and will be referred as *O. formosanus* as previously suggested^18,22^.

Analyses of historical demographic trajectories under a pairwise sequentially Markovian coalescent (PSMC) model indicated that all lineages experienced demographic expansions approximately 139,900– 64,600 years ago, prior to the Last Glacial Maximum (Fig. 1e). Peak effective population sizes (*N_e_*) varied considerably, with Nagano Masu (∼381,000) and Qijiawan (∼375,000) having the largest historical *N_e_*. However, all populations underwent pronounced contractions during and post glaciation, particularly severe in the Taiwanese populations, resulting in recent *N_e_* below 28,000 and 34,000 individuals for Qijiawan and Hehuan creeks, respectively. We further calculated the relative proportions of homozygous and heterozygous variants in both synonymous and loss-of-function (LOF) mutations. Across all populations, heterozygous derived LOF variants were more frequent than homozygous derived LOF variants (Fig S7), consistent with the expectations that smaller populations would accumulate higher frequencies of deleterious alleles and increased homozygosity. In particular, the Qijiawan creek population showed a significantly higher proportion of both heterozygous and homozygous derived LOF variants compared to synonymous variants, highlighting its heightened genetic vulnerability (Fig. S7). Together, these findings emphasize the profound evolutionary distinctiveness and genetic vulnerability of *O. formosanus* populations.

### Extensive chromosome fusions and sex chromosome rearrangements in *O. formosanus*

Genome-wide synteny analyses uncovered substantial chromosomal rearrangements differentiating masu salmon lineages, using rainbow trout (*O. mykiss*), *O. keta* and *S. fontinalis* as outgroups (Fig. 2a and Table S6). Eight chromosomes of the masu salmon complex originated exclusively from fusion events involving 16 ancestral chromosomes identified through comparison with homologous chromosomes in *O. mykiss*, reflecting considerable genomic reorganization since their divergence (Fig. 2a and Table S6). Notably, *O. formosanus* underwent further chromosomal reorganizations through telomere-to-telomere fusion events involving at least 14 ancestral chromosomes, in contrast to only two in *O. m. masou* (Fig. 2a and Table S6). For example, chromosome 1 of *O. formosanus* corresponds to chromosomes 23 and 24 in *O. m. masou* and chromosomes 5, 29, 1, and 9 in *O. mykiss*, while chromosome 4 corresponds to chromosomes 4 and 5 in *O. m. masou* and chromosome 17 in *O. mykiss*, illustrating significant ancestral chromosome fusions. These fusion sites are enriched in repeats including satellite DNA, characteristic of centromeric repositioning or neocentromere formation^41^. In addition, genes residing at these *O. formosamous* chromosomes displayed significantly elevated evolutionary rates compared to non-fused chromosomes (dN/dS ratios, P=0.009, Wilcoxon Rank Sum test, Fig. S8), suggesting that structural genomic rearrangements facilitated adaptive evolution in response to ecological pressures such as cold tolerance and metabolic specialization^42,43^.

**Figure 2.**
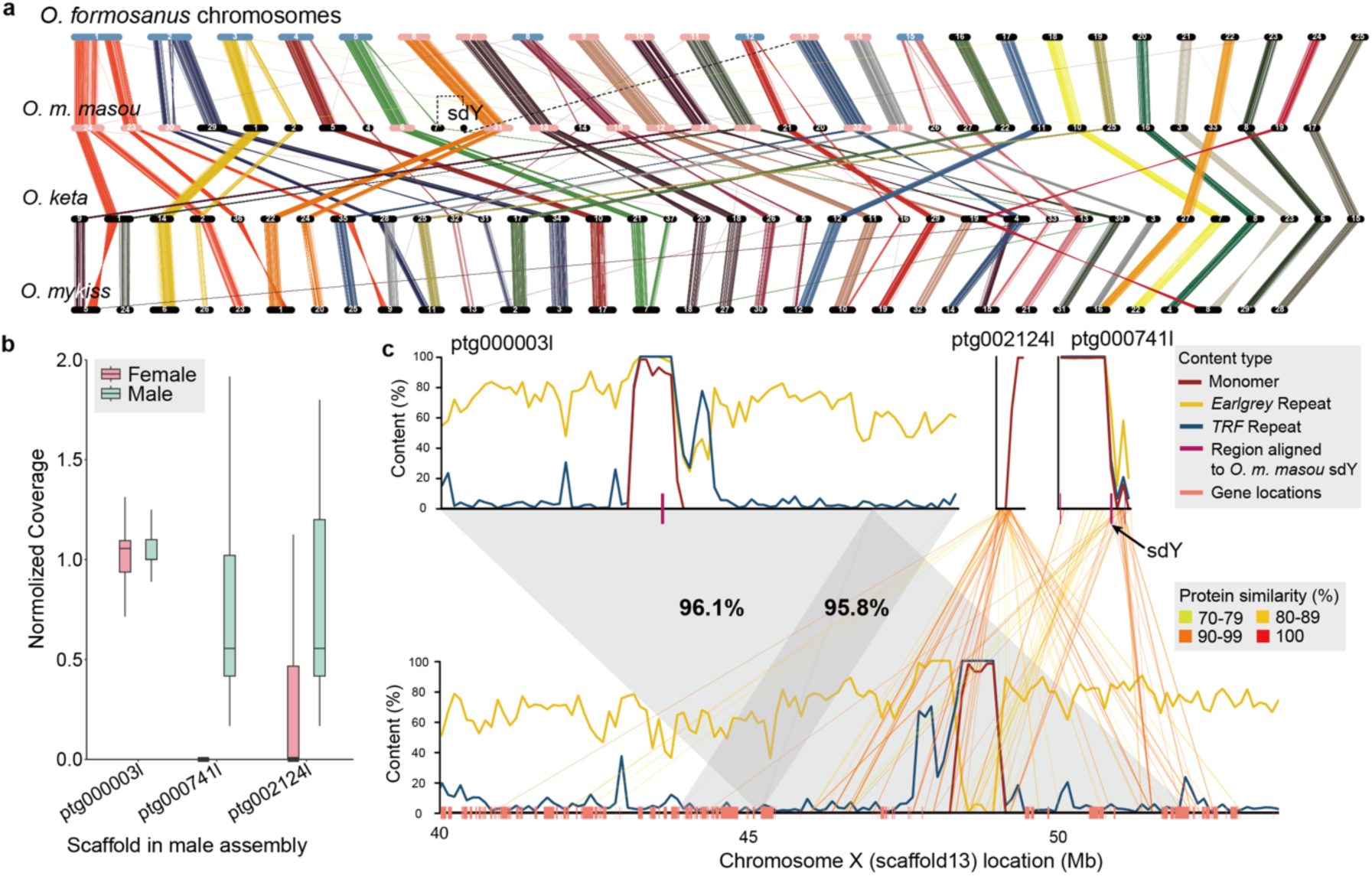
Chromosome synteny and sex chromosome evolution in *O. formosamous*. (**a**) Genome-wide synteny comparison across *Oncorhynchus* species. Chromosomes highlighted in blue denote fusions unique to *O. formosanus*, and those in red indicate fusions in the common ancestor of masu salmon lineages. Colored lines represent pairwise single-copy orthologs. The scaffold containing the sex-determining gene (*sdY*) in *O. m. masou* follows data from Christensen et al. ^28^. (**a**) Normalized coverage of female (pink) and male (blue) sequencing reads mapped onto selected scaffolds from the male *O. formosanus* genome assembly. (**c**) Detailed annotation of repetitive elements and sequence alignments between putative X-linked (ptg00003l) and Y-specific scaffolds (ptg000741l and ptg002124l) from the male genome, aligned to chromosome X (scaffold13) from the female assembly. Numbers above shaded grey area indicate nucleotide identity between X chromosome sequences of male and female assemblies. Lines connecting Y-specific scaffolds (ptg000741l and ptg002124l) and scaffold13 represent homologous regions, with colors indicating amino acid sequence similarity. The exact location of the *sdY* gene on scaffold ptg000741l is indicated.

The sex-linked region demonstrated particularly pronounced chromosomal rearrangements. An earlier study linked the contig containing the sex-determining gene (sdY) to chromosome 7 in *O. m. masou*, although alternative arrangements were suggested for the Amago and Biwa lineages^28^. Our genome assembly originated from a single female revealed that this *O. m. masou* contig was aligned uniquely to the central region (∼50 Mb) of chromosome 13 (Fig. S9). This location corresponds exactly to a known ancient chromosome fusion site homologous to chromosomes 14 and 25 in rainbow trout (*O. mykiss*) (Fig. 2a and Fig. S10) and contains abundant 281-mer satellite repeats typical of centromeric regions.

To further clarify this genomic region, we produced an additional genome assembly from a male individual (Methods). Three scaffolds were mapped uniquely to this fusion region and revealed clear coverage differences between males and females, enabling us to unambiguously distinguish XY-shared contigs from Y-specific ones (Fig. 2b). An inversion was identified between the XY-shared region between the two assemblies, and the sdY gene was identified specifically on a Y-specific contig, closely adjacent to the Y chromosome’s centromeric region (Fig. 2c). Interestingly, sdY is also located ∼ 5Mb adjacent to the centromere of *O. mykiss* non-homologous Chr29, consistent with sdY’s mobility in fishes^44,45^. Pairwise identity comparisons between homologous X and Y-specific regions from these contigs revealed elevated sequence divergence averaging 9.6%, indicating that XY divergence was restricted to the genomic segment spanning from sdY to the centromere, a pattern similarly observed in rainbow trouts^44^. Our findings demonstrate significant differences in sex chromosome configurations within the *O. masou* complex, highlighting how chromosomal rearrangements have shaped genomic evolution and potentially contributed to reproductive isolation.

### Proteome specialization of *O. formosanus*

We investigated gene and protein domain expansions specific to *O. formosanus* by annotating protein family (Pfam) domains across salmonid genomes (Fig. S11a). Several Pfam domains exhibited significant expansions, notably the immune-related C1-set immunoglobulin domain (>60 copies), chromatin-associated Chromo domain (∼60 copies), glycosyltransferase (Glyco_transf_49; ∼60 copies), and glycoside hydrolase (Glyco_hydro_56; ∼13 copies). Interestingly, we identified an enrichment of microtubule-associated protein MAP6 (∼30 copies), which are known to confer cold stability to microtubules^46^ and upregulated during acute cold stress in Japanese flounder^47^. Aligning with this cold-adaptive expansion, we observed a substantial enrichment of the ice-binding lectin domain (LbR_Ice_bind; ∼19 copies) in *O. formosanus*. Ice-binding proteins (IBPs) protect cells by binding to and inhibiting ice crystal growth. Our phylogenetic analyses indicated that these expanded LbR_Ice_bind proteins in *O. formosanus* formed a lineage distinct from previously characterized ice-structuring proteins (ISP types I, II, and IV^48^) identified in other *Oncorhynchus* species (Table S7 and Fig. S12), suggesting coordinated functional specialization to colder freshwater habitats. Orthologous group expansions via CAFE analyses further highlighted significant enrichment in 331 genes associated with exocrine and salivary gland development, chromatin assembly, DNA conformation changes, homophilic cell adhesion, innate immune responses, and calcium ion transport (Table S8), underpinning key physiological adaptations to freshwater environments. In contrast, closely related *O. m. masou* exhibited distinctly different expanded domains (e.g., Beach and Polycystin domains, Fig. S11b), reflecting divergent evolutionary trajectories post-geographic separation^49^.

### Divergent population structures among *O. formosanus* populations

To resolve fine-scale genetic structure and evolutionary differentiation within *O. formosanus*, we analyzed 1,509 SNPs derived from whole-genome sequencing and ddRADseq datasets (Table S1 and Fig. S13), applying a 1-Mb window selection approach across individuals from the three major stream systems. Principal component analysis (PCA) revealed distinct clustering patterns that mirrored geographic distributions (Fig. 3a). Importantly, populations from the Hehuan creek (HEH) exhibited substantially greater dispersion along both principal component axes compared to the other groups, occupying an expanded region in multivariate space. In contrast, Luoyewei creek populations and the downstream sites of Hehuan creek (HEH_01, HEH_02) formed tightly clustered, discrete genetic groups, suggestive of localized genetic homogenization and elevated genetic diversity within the Hehuan creek.

**Figure 3.**
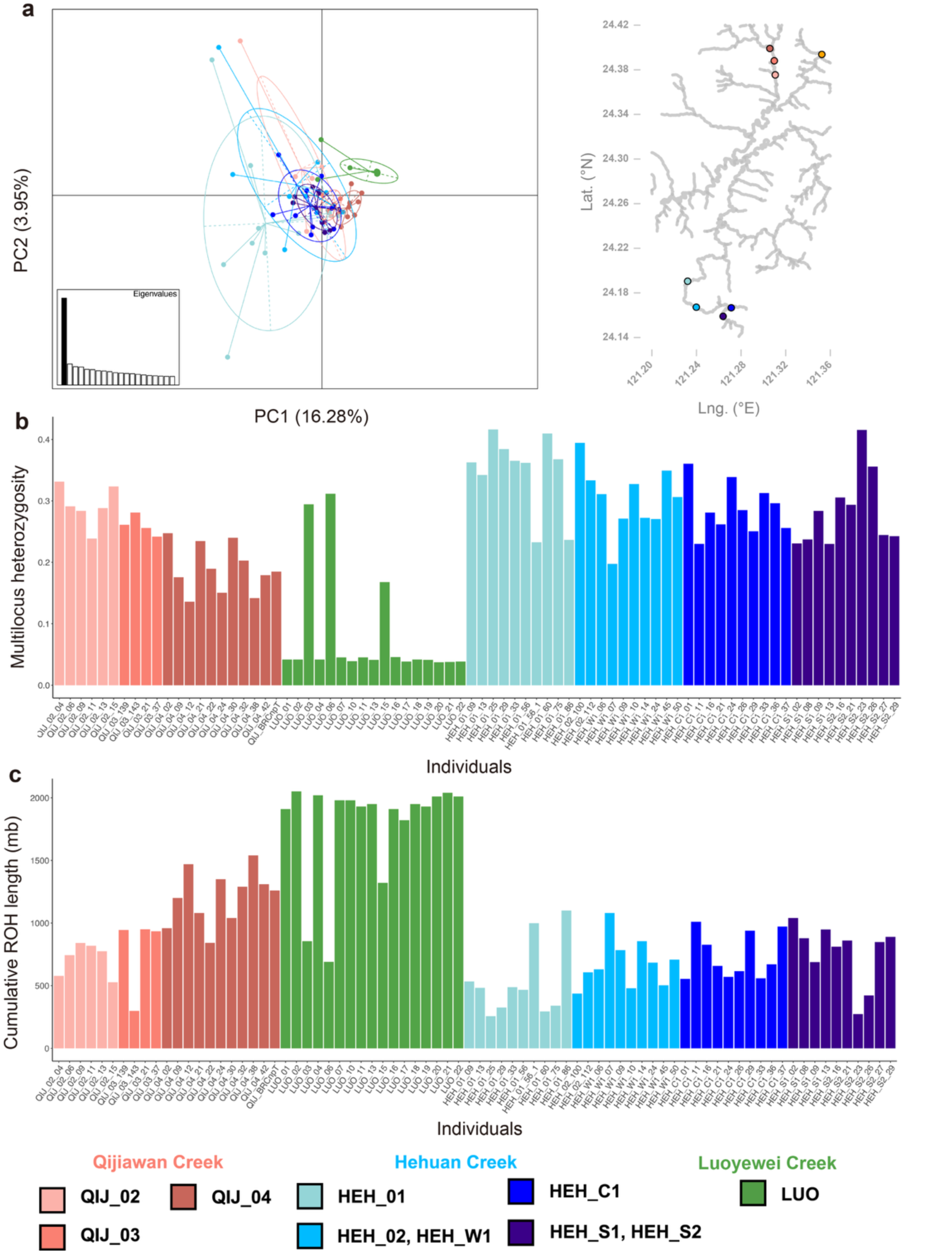
Population structure, genetic diversity and level of inbreeding of the Formosan salmon. (**a**) PCA result of the population structure shows no significant differentiation among most populations, except for those from Luoyewei Creek and the downstream sites of Hehuan Creek (HEH_01, HEH_02, HEH_W1). Populations from Hehuan Creek occupy a broader range in PCA space (PC1 vs. PC2), indicating a higher genetic diversity. (**b**) Multilocus heterozygosity across individuals from the three creek systems. Hehuan Creek populations exhibit the highest genetic diversity, followed by Qijiawan Creek and lastly by Luoyewei Creek. (**c**) The level of inbreeding, estimated using runs of homozygosity (ROHs), is highest in Luoyewei Creek, intermediate in Qijiawan Creek, and lowest in Hehuan Creek.

### Long-term native status of the Hehuan lineage

To further investigate if the Hehuan creek system harbor potential, previously unrecognized reservoir of genetic variation within *O. formosamous*, we assessed multilocus heterozygosity (MLH) and runs of homozygosity (ROH) across individuals. Hehuan creek samples consistently exhibited the highest heterozygosity and lowest levels of inbreeding among all creeks (MLH: HEH–QIJ mean difference = 0.07, P < 0.05; HEH–LUO = 0.23, P < 0.05; ROH: HEH–QIJ = –310,968.3, P < 0.05; HEH–LUO = –1,108,330, P < 0.05) (Fig. 3b–c), followed by Qijiawan creek (MLH: QIJ–LUO = 0.15, P < 0.05) and lower inbreeding levels (ROH: QIJ–LUO = –797,361.8, P < 0.05), while Luoyewei creek samples showed markedly reduced heterozygosity and elevated inbreeding. The ROH patterns in Hehuan individuals—characterized by fewer and shorter homozygous segments— are indicative of demographic continuity and argue against a recent or artificially-introduced origin. Within Qijiawan creek, we also detected a clear longitudinal gradient in genetic diversity (Pearson’s r: −0.82, P<0.05): downstream populations retained higher heterozygosity and lower inbreeding compared to upstream ones. This pattern aligns with theoretical expectations from metapopulation dynamics, wherein downstream connectivity supports gene flow while upstream isolation fosters drift^50^. In contrast, the Luoyewei creek showed no such spatial structure, consistent with its known history of hatchery stocking and limited founder input.

### Demographic patterns across stream populations

Population genetic analyses revealed distinct demographic parameters across streams. Effective population size (*N_e_*) measurements showed significant differences among populations: Luoyewei Creek had *N_e_* <1, while Hehuan and Qijiawan populations maintained substantially higher values of 98.5 and 84.5, respectively (Fig. 4a). Temporal reconstruction analyses of historical *N_e_* revealed divergent trajectories (Fig. 4b). The data showed parallel demographic patterns between Hehuan and Qijiawan until approximately 100 years ago, when they began to diverge coinciding with the 1930s conservation designation of Qijiawan Creek populations as a natural monument during the Japanese administration. Both populations showed marked declines beginning in the 1960s, corresponding with the construction of the Central Cross-Island Highway. Our *N_e_* reconstruction from Fig. 4b showed different recovery patterns in effective population size: Qijiawan Creek has exhibited gradual recovery over the past two decades, while Hehuan Creek’s *N_e_* increased notably following the 2017 reintroduction program^24,25^. Critically, post-2017 genomic data from Hehuan Creek revealed higher-than-expected genetic diversity—reflected in elevated multilocus heterozygosity (MLH), reduced runs of homozygosity (ROH), and the presence of unique allelic variants not found in the Qijiawan source population (Fig. 3b–c, Table S1). Additionally, population structure analyses identified a discrete genetic cluster within Hehuan Creek, suggesting the persistence of a previously undetected native lineage, now likely admixed with reintroduced individuals.

**Figure 4.**
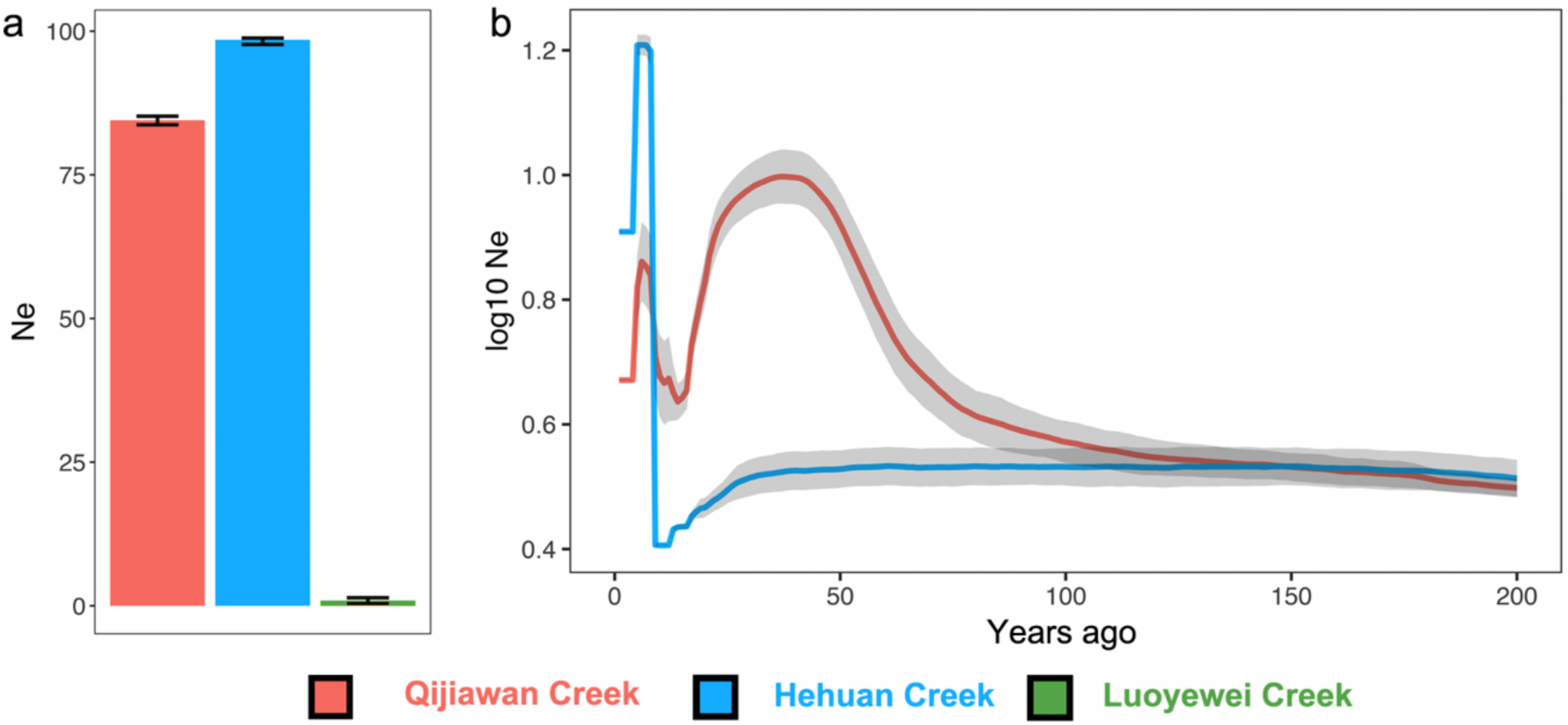
Current effective population size *N_e_* and demographic history of Formosan salmon populations. (**a**) Estimated current *N_e_* for Formosan salmon populations across multiple stream sites. The Qijiawan and Hehuan populations appear to maintain effective sizes above thresholds associated with inbreeding depression, while the Luoyewei Creek population, which likely comprises individuals from hatchery releases, has a very small effective population size. (**b**) Inferred demographic histories of populations from Qijiawan and Hehuan Creeks. The contrasting trajectories reflect differences in conservation management and habitat development across the two regions.

### Life table analyses reveal stream-specific demographic asymmetries and typhoon-induced shifts

Using improved non-lethal sampling methods and bootstrapped statistical inference, we generated age-structured life table models for *O. formosanus* populations in the two headwater stream systems (Methods). These estimates revealed significant impacts of typhoon on population dynamics and important differences between stream systems. Under non-typhoon scenarios, both streams exhibited strong intrinsic growth potential (dominant eigenvalue λ > 1.0), with QIJ at λ = 2.23 (1.53 – 2.65) and HEH at λ = 2.25(1.84 – 2.83), indicating rapid population growth capacity. However, under typhoon disturbance scenarios, λ values dropped substantially below 1 (QIJ = 0.405 (0.01-0.55); HEH = 0.477 (0.26-0.66)), indicating population contraction (Table S9). Notably, HEH maintained a relatively higher growth rate post-typhoon, demonstrating greater resistance to disturbance. Survival rate analysis showed that annual survival during the typhoon period (January→November) was markedly lower than during baseline periods (November→June) (Table S10). Under typhoon influence, annual survival rates for age-0 fish in QIJ and HEH were 0.202 and 0.184, respectively, while under baseline conditions they were 0.616 and 0.615. Adult fish (ages 1-2) exhibited reduced survival rates during typhoon periods (0.046-0.131), indicating that typhoons imposed substantial survival pressure across all age classes. Recruitment analysis revealed significant declines in recruitment capacity due to typhoons. In January 2024 (non-typhoon), adjusted recruitment for QIJ and HEH was 1,492 and 2,354 individuals/1000m², respectively, but decreased to 657 and 634 individuals/1000m² in January 2025 (post-typhoon), representing declines of 56-74%. Fecundity coefficients were similarly affected (Table S11), with maximum fecundity under non-typhoon scenarios reaching 19.3 and 8.9 for QIJ and HEH, respectively, while post-typhoon values declined to 1.53 and 1.21, indicating substantial reductions in reproductive capacity. Overall, these life table estimates demonstrate that typhoon disturbance causes comprehensive negative impacts on *O. formosanus* populations, including concurrent declines in survival, recruitment, and fecundity. HEH retained higher post-disturbance λ, exhibited superior resistance to disturbance compared to QIJ, a difference that has important implications for conservation strategy development under climate change.

### Divergent long-term responses of Qijiawan and Hehuan populations to typhoon disturbances

Based on empirically derived demographic parameters—including age-specific survival rates, recruitment levels, and fecundity estimates corrected for sampling bias—we developed a stochastic, discrete-time simulation model to assess the long-term population dynamics of *O. formosanus* under varying typhoon disturbance regimes. The model incorporates a two-stage Leslie matrix structure representing pre-disturbance (January–November) and non-disturbance (November–June) periods, with transitions modulated by observed survival patterns (_1−11_ and 𝑆_11−6_; Table S10). Simulation results revealed divergent trajectories and extinction probabilities between the QIJ and HEH systems under varying typhoon disturbance frequencies (Fig. 5a–d). At low disturbance (p = 0.3) both populations stabilised after an initial spike, but at different plateaus: HEH settled at ≈ 1,100–1,150 fish · 10³ m⁻² whereas QIJ levelled off at ≈ 1,000– 1,050 fish · 10³ m⁻². When p rose to 0.5, HEH maintained a gently declining yet robust abundance of ∼550–650 individuals, while QIJ declined steadily from ≈ 600 to ≈ 450 individuals over 20 years. Under high disturbance (p = 0.7) both curves plunged, yet HEH fell from ∼600 to ∼50 individuals—hovering just above the quasi-extinction threshold—whereas QIJ dropped from ∼450 to < 40 individuals, crossing the threshold in most replicates. At p = 0.9 both streams trended toward local extinction: QIJ fell below 10 individuals by year 8 and approached zero thereafter, while HEH slipped below the threshold by year 9 and dwindled to < 5 individuals by the simulation endpoint.

**Figure 5.**
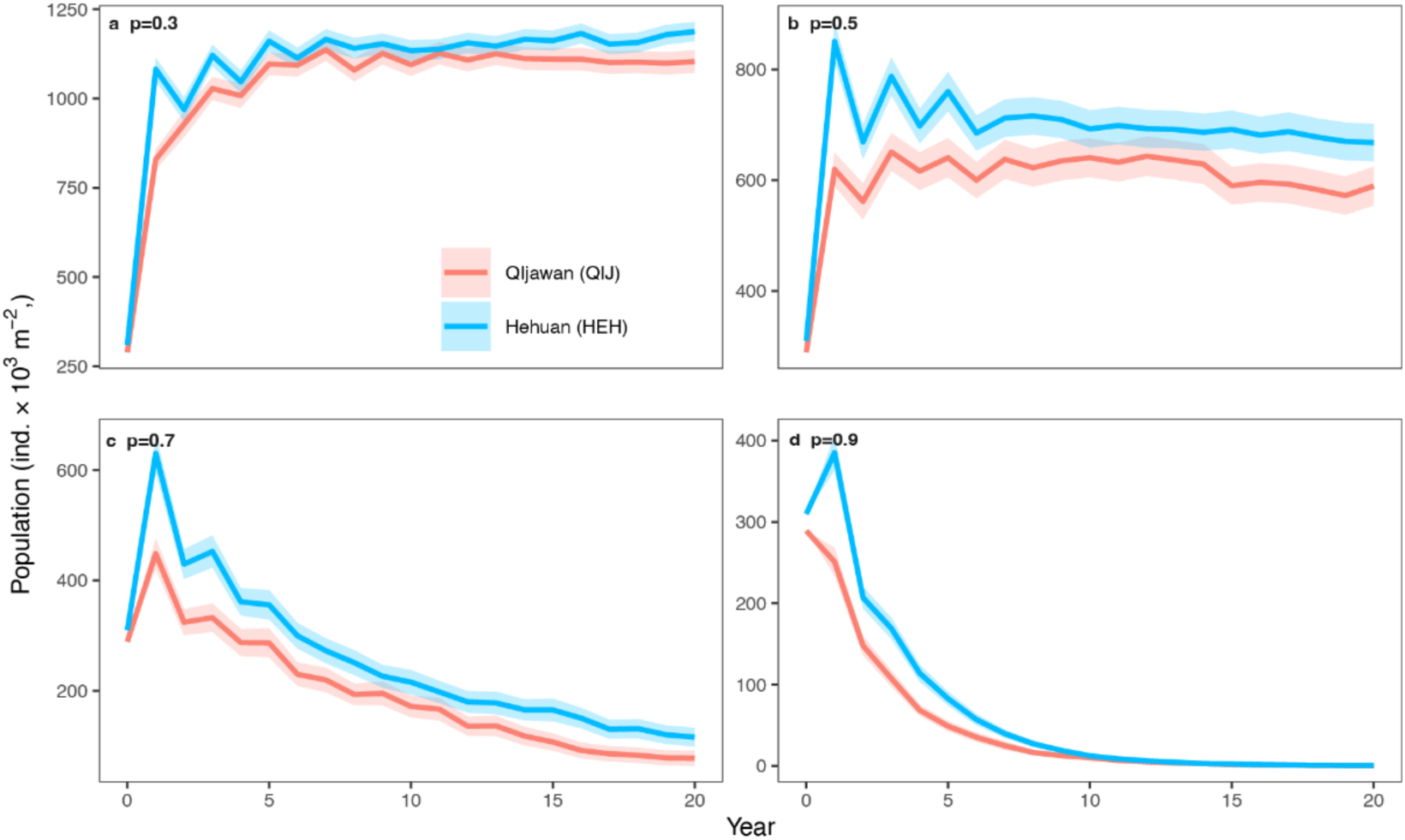
Population trajectories and extinction risk under varying typhoon frequencies. (a-d) Simulated population dynamics over 20 years under different annual typhoon occurrence probabilities (p = 0.3, 0.5, 0.7, 0.9) over a 20-year simulation period. Panels a–d represent different disturbance scenarios, with each line indicating mean abundance (individuals aged 1-4, per 1,000 m²) for Qijiawan (QIJ; red) and Hehuan (HEH; blue) stream populations. Shaded bands represent 95% confidence intervals across 1,000 stochastic replicates per scenario. Simulations incorporated stage-specific survival rates, Leslie matrix projections, and probabilistic typhoon impact modeled using zero-inflated Gamma distributions derived from historical records.

Extinctionrisk analysis using a quasiextinction threshold of ≤ 10 individuals per 10³ m² corroborated these trends (< 2 mature female per 10³ m²). Across 1,000 replicates, probabilities of QIJ climbed from 2.2 % (95 % CI 1.4–3.3 %) at p = 0.3 to 29.8 % (27.0–32.7 %) at p = 0.5, 85.7 % (83.4–87.8 %) at p = 0.7, and 100 % (99.6–100 %) at p = 0.9. The corresponding risks for HEH were 0.5 %, 15.1 %, 77.4 %, and 99.9 %. Fisher’s exact tests yielded odds ratios (QIJ / HEH) of 4.48 (1.69–11.86) at p = 0.3, 2.39 (1.92–2.97) at p = 0.5, and 1.75 (1.39–2.20) at p = 0.7, confirming that QIJ remains 1.7 to 4.5 times more vulnerable until both populations approach certain extinction (Fig. S14). LogisticGLM contrasts showed a monotonic decline in OR with increasing disturbance, reflecting convergence of relative risk as absolute probabilities approach unity. These results indicate that while both systems are vulnerable to high-frequency typhoon disturbance, QIJ exhibits substantially greater extinction risk across all disturbance scenarios.

## Discussion

### Adaptive capacity and conservation implications in marginal populations

Our findings reveal how small, geographically marginal populations can maintain adaptive potential under prolonged environmental constraint. *Oncorhynchus formosanus*, now confined to a few high-mountain streams in Taiwan, represents the southernmost lineage of the genus and is not a recent derivative of Japanese masu salmon. Phylogenomic analyses of 1,484 single-copy nuclear genes place its divergence from *O. m. masou* over one million years ago during the Pliocene epoch. In contrast, ddRAD-based coalescent modeling suggests that the last gene flow between Taiwanese populations and their southern Japanese relatives (Nagano Masu and Amago) occurred around 50,000 years ago. These distinct timelines reflect different evolutionary layers—deep ancestral divergence versus more recent lineage sorting—and arise from the differing temporal sensitivity of genomic markers. This ancient divergence is supported by genealogical divergence index (GDI) values exceeding established species-level thresholds^39,40,51^. We also observed substantial chromosomal fusions accompanied by elevated dN/dS ratios, along with notable expansions in gene families associated with cold adaptation, such as *MAP6* and *LbR_Ice_bind*. These adaptive signatures have persisted despite restricted gene flow and historical bottlenecks, likely facilitated by Taiwan’s complex montane river systems that both isolate populations and maintain localized connectivity. Such genomic plasticity, enabled by historical whole-genome duplications (WGD), is also documented in Atlantic salmon, where duplicated genomic regions provide raw material for rapid adaptation to local environments^31^. Similar patterns of local adaptation, where native populations outperform non-native ones, have been widely reported across salmonids^6^. Our extensive multi-stream sampling revealed genetically distinct *O. formosanus* populations with diversity levels that rival, and in some cases exceed, those of Japanese counterparts. However, effective population size (*N_e_*) estimates show stark contrasts: Luoyewei populations exhibit extremely low *N_e_* (<1), consistent with their hatchery origin, while Hehuan and Qijiawan creeks maintain higher values (98.5 and 84.5, respectively). These estimates exceed the threshold (*N_e_* < 50) needed to minimize short-term inbreeding depression^52,53^, but still fall within the range (50 < *N_e_* < 500) that warrants targeted conservation intervention.

### Historical conservation impacts and demographic trajectories

The contrasting trajectories of the Hehuan (HEH) and Qijiawan (QIJ) populations illustrate how the timing, enforcement, and ecological targeting of conservation measures shape long-term demographic outcomes. Until approximately a century ago, both lineages apparently followed similar patterns. Their histories began to diverge in the 1930s when *O. formosanus* was designated a natural monument under the Japanese colonial rule^54,55^. Restrictions on harvest and agricultural activity were implemented across the upper Dajia River basin, and these measures appear to have been more effectively enforced in Qijiawan, contributing to early signs of demographic recovery. In contrast, similar bans in Hehuan Creek may have had limited impact due to the remoteness and inaccessibility of the area at the time.

Despite these early interventions, both populations experienced sharp declines in the 1960s, coinciding with the construction of the Central Cross-Island Highway and widespread habitat fragmentation^56^. These patterns underscore that *O. formosanus* retains recovery potential when habitats receive sustained and effective protection^57^. However, the trajectory differences between HEH and QIJ in recent decades suggest that early legal protections alone are insufficient without continued ecological management tailored to each stream’s conditions.

Our demographic simulations (Fig. 5) support this view. Under increasing typhoon disturbance probabilities, QIJ populations declined steadily under all disturbance levels, whereas HEH populations maintained higher abundance and slower decline, remaining stable or slightly increasing only when typhoon frequency was low-to-moderate (p ≤ 0.5). These trends align with survival estimates across age classes, and with HEH’s more favorable age structure and fecundity profile. The observed divergence highlights the critical role of ongoing, stream-specific ecological support—such as habitat quality and spawning condition maintenance—in sustaining viable populations.

While genetic rescue might appear attractive for isolated or declining populations, our results caution against indiscriminate translocation in deeply diverged lineages. Introducing genetic material from non-local sources can lead to hybrid incompatibility, disruption of co-adapted gene complexes, and genomic swamping^58,59^. The case of the Chinese giant salamanders, where unregulated stocking erased native lineages through uncontrolled admixture^60^, exemplifies these risks. Thus, any intervention must be informed by both genomic distinctiveness and local demographic context to avoid undermining evolutionary legacy under the guise of recovery.

### Rediscovery of native lineages and stream-specific recovery dynamics

Our findings reveal that population recovery in *O. formosanus* varies significantly across its range, with genetic composition differing markedly among streams. Most striking is the case of Hehuan Creek, previously assumed devoid of native salmon since the 1960s due to overharvesting^56^. Contrary to this assumption, our whole-genome sequencing uncovered a genetically distinct lineage consistent with a surviving native population. This discovery coincided with a reintroduction program initiated in 2017^25^, which contributed to Hehuan’s population increase. However, genomic signals—elevated heterozygosity, reduced inbreeding coefficients, and unique allelic variants—indicate that this growth cannot be attributed to stocking alone. Instead, our data suggest a synergistic scenario where reintroduced individuals may have encountered and interbred with remnant natives under favorable ecological conditions. This interpretation is supported by the recent study^24^, which demonstrated that Hehuan Creek provides superior environmental conditions for salmon reproduction compared to the long-protected Qijiawan Creek. Similarly, Qijiawan populations have shown demographic recovery over the past two decades, aligning with intensified conservation measures including habitat restoration and regulated fishing. In stark contrast, Luoyewei Creek populations, despite similar census-based recovery, display extremely low genetic diversity and no evidence of increasing effective population size. This discrepancy highlights a key limitation of conventional conservation aquaculture: numerical recovery alone is insufficient to ensure evolutionary viability. As demonstrated in other systems^61,62^, hatchery programs relying solely on long-term broodstock often fail to maintain genetic diversity, leading to elevated inbreeding and loss of rare alleles. In comparison, approaches based on wild-origin offspring collection consistently preserve broader genetic variation and more accurately represent natural spawner diversity. The Luoyewei case mirrors such outcomes, where demographic recovery masks underlying genetic erosion due to historical reliance on hatchery stock with limited founders. Without revisiting propagation protocols to incorporate alternative strategies—such as wild-origin embryo collection, sibship-based pedigree reconstruction, or genomic monitoring—such populations may continue to erode in adaptive potential despite apparent demographic recovery. These findings underscore the importance of embedding genetic resilience as an explicit conservation goal, particularly in isolated headwater systems where natural recolonization is unlikely and demographic indicators alone may be misleading.

### Integrating genomics with ecological context for effective conservation

The differing responses observed among the three stream systems suggest that traditional, uniform conservation approaches may have important limitations when applied across ecologically diverse contexts. Our demographic simulations reinforce this point: under simulated typhoon disturbances, HEH shows markedly greater persistence, but still declines under high-frequency events (p ≥ 0.7), whereas QIJ collapses under every scenario studied (Fig. 5c–d). These divergent outcomes likely stem from differences in genetic diversity, local adaptation, and life-history traits shaped by stream-specific ecological conditions—not simply by broad variables like elevation or streamflow. Qijiawan’s vulnerability under high disturbance regimes stems not only from reduced genetic variability, but also from demographic constraints: lower juvenile survival, decreased fecundity, and narrower terminal age class representation. These factors collectively reduce its ability to recover following extreme events, as clearly observed in the modeled outcomes. As such, ecological context must inform genetic interpretations— demographic plasticity and reproductive capacity are critical buffers that genomic data alone cannot predict. Such findings are part of a broader pattern repeatedly observed in salmonids: local adaptation is both common and consequential. Numerous studies have shown that even fine-scale environmental variation can drive significant divergence in traits critical for survival and reproduction. For example, Fraser River Sockeye Salmon populations facing more challenging migration routes exhibit enhanced aerobic scope and cardiac performance^63^, demonstrating physiological traits adapted to local demands. Similarly, thermal physiology in Pacific salmonids varies widely among populations, reinforcing the need for context-dependent conservation approaches rather than generalized prescriptions^64^.

Hehuan Creek provides a compelling case: despite historical overharvesting, it now supports a genetically distinct and resilient population. Genomic data reveal elevated heterozygosity, low inbreeding, and unique allelic variants—signals not fully explained by recent reintroduction efforts^25^. Instead, these patterns point toward a hybrid population formed through interaction between remnant natives and reintroduced fish, thriving in an ecologically favorable habitat^24^. This underscores the importance of integrating ecological niche data with genomic signals to avoid underestimating the recovery potential of marginal populations. Yet, local adaptation also complicates well-intentioned interventions. A recent reciprocal transplant experiment in brown trout populations by Labonne et al.^65^ found little evidence of genetic rescue and no consistent fitness advantages for foreign or hybrid individuals, suggesting that gene flow alone does not guarantee improved outcomes. These results caution against assuming that translocations or supplementation will necessarily benefit recipient populations, especially when local selection regimes are strong or not fully understood. Taken together, these findings argue for conservation strategies that are highly tailored, genetically informed, and ecologically grounded. Preserving intraspecific diversity requires more than demographic stabilization; it demands recognition of local evolutionary trajectories and the complex interplay between genes, environment, and behavior.

## Materials and Methods

### Sample Collection

For whole genome sequencing, a female adult (OIJ-BRCnpT, **Table S1**) and a female subadult (CHI-04-2409-01, **Table S1**) of Taiwanese salmon, *O. formosanus*, were collected from the Qijiawan Creek (24°23’58.5"N 121°18’19.0"E), Heping District, Taichung City, Taiwan. The brain, eye, gill and odor tissues were excised from fish body and preserved in liquid nitrogen or RNAlater™ Stabilization Solution (Cat: AM7021, Invitrogen, Thermo Fisher Scientific Inc.) at −80°C freezer. The muscle tissues were preserved in 100% ethanol or liquid nitrogen at −80°C freezer.

To investigate nucleotide diversity and population genetics of Taiwanese salmon, each fresh fin tissue of 44 Taiwanese, three Japanese amago and two Japanese Nagano Masu individuals was collected and sequenced via ddRAD-seq (Table S1 and Fig. S1). Each captured salmon was anaesthetized with 2-phenoxyethanol and immediately released to the original habitats after fin clipping. Six of 44 Taiwanese individuals (QIJ_02_04-, QIJ_02_15, LUO_07, HEH_01_13, HEH_W1_24, HEH_02_112, QIJ-BRCnpT) and five Japanese samples were further selected for Illumina PE-150 sequencing. All experiments in this study were performed in accordance with guidelines of the animal ethics committee and were approved by the Institutional Animal Care and Use Committee, Academia Sinica (IACUC 24-09-2284) and collection permissions of Ministry of Agriculture (MOA) and National Park Service (NPS), Ministry of the Interior.

### DNA and RNA sequencing

For genome sequencing, 30-50 mg of muscle tissue was lysed with proteinase K and Biofluid & Solid Tissue Buffer from Quick-DNA™ HMW MagBead Kit (Catalog No. D6060, Zymo Research Corp., USA), following the manufacturer’s instructions. For ddRADseq, 2-10 mg of fin tissue was lysed with proteinase K and BL2 Buffer from MagPurix Tissue DNA Extraction Kit (Catalog No. ZP02004, Zinexts Life Science Corp., Taiwan), following the manufacturer’s instructions. The eluted genomic DNA were stored at −20°C before sequencing.

For PacBio long-read sequencing, the genomic DNA was enriched for fragments larger than 10 kb using the Short Reads Eliminator Kit (PacBio). DNA purity was checked using SimpliNanoTM - Biochrom Spectrophotometers (Biochrom, MA, USA). DNA concentration was determined by Qubit 4.0 fluorometer (Thermo Scientific) and the fragment size was monitored by the Qsep 100TM system (Taiwan). The SMRTbell library was prepared according to the Preparing whole genome and metagenome libraries using SMRTbell® Prep Kit 3.0 procedure (PacBio). In brief, 1 ug of high molecular weight genomic DNA was digested with DNase to remove single-strand overhangs, following end repaired and added dA-tail using KAPA End Repair and A-tailing reagent (KAP HyperPlus Kit, KAPA Biosystems, Roche). The Overhang Adapter was introduced onto double-stranded DNA fragments by ligation steps. After purification with AMPure PB beads to remove the adapter dimer, the SMRTbell library was incubated with sequel II Binding Kit 3.2 for the primer annealing and polymerase binding. At last, sequencing was performed in the HiFi sequencing mode on a PacBio Sequel IIe instrument to generate the subread.bam raw data for 30 hours movie time. It was then sequenced through the services of the BIOTOOLS Co., Ltd. and the High Throughput Genomics Core Facility at Academia Sinica, Taiwan.

For Illumina short-read sequencing, genomic DNA was sheared using a Covaris 2 machine, and then quantified using a Qubit 2.0 Fluorometer. A short-read sequencing library was also constructed from 200ng of the sheared gDNA using the TruSeq Nano DNA High Throughput Library Prep Kit (Cat. No. 20015965). The PE-150 reads were then sequenced using an Illumina NovaSeq X Plus through the Genomics BioScience & Technology (New Taipei City, Taiwan). In addition, the high-resolution chromosome conformation capture (Hi-C) libraries were prepared and sequenced using Illumina NextSeq 2000 at the High Throughput Genomics Core Facility at Academia Sinica, Taiwan.

For Illumina RNA sequencing, five to 30 mg of brain, eye, gill and odor tissues were excised and homogenized with 600 µl Trizol solution for RNA sequencing. RNA was extracted from the homogenized Trizol solution using RNeasy® Plus Mini Kit (Cat: 74134, Qiagen), and quantified using the Agilent 2100 Bioanalyzer. A short-read sequencing library was constructed from ∼1.5μg of the quantified RNA using the TruSeq Stranded mRNA Library Prep Kit (Cat. No. 20020595). It was then sequenced on the Illumina Novaseq X Plus through the services of GENOMICS BIOSCIENCE & TECHNOLOGY CO., LTD, Taipei, Taiwan.

### Male-specific marker detection

For sex determination, polymerase chain reaction (PCR) using male-specific primer pair OtY2m (5’-TCAATCTGTGACGTCACTCA-3’ and rev: 5’-GGCTTACCGCTCCCAAGTAT-3’) ^66^ were conducted on 50-100 ng of genomic DNA in 25-μl reactions from each individual of the *O. masou* complex with the following composition: 12.5 μl of GoTaq G2 Green Master Mix (M7822, Promega), 2 μl of each OtY2m (5μM, forward and reverse) primer, 6.5μl of dd water and 2μl of genomic DNA. The amplification program was 94 °C for 2 min; followed by 35 cycles of 94 °C for 1 min, 52 °C for 1 min, and 72 °C for 1 min; followed by 72 °C for 2 min. The final products were analyzed using 2% agarose gel for 30-min electrophoresis and fragment visualization.

### Genome assembly and gene annotation

Hifiasm (ver 0.19.9)^67^ was used to assemble the raw PacBio Hifi reads and haplotigs were removed using HaploMerger2 (ver. 20180603) ^68^. The assembly were further scaffolded using HiC reads with YaHS ^27^ and subsequently curated in Juicebox^69^. Repetitive DNA sequences were identified and masked using EarlGrey^70^. The proteomes of 11 Salmons were obtained from NCBI (listed in **Table S2**). Transcriptome reads of eye, brain, gill and odor were mapped to the genome assemblies using STAR (ver. 2.7.7a) ^71^. The gene models were predicted using BRAKER3 (ver 3.0.8)^72^ with proteomes and RNA-seq mappings as evidence hints. Protein families and GO terms were annotated using Pfam scan (https://ftp.ebi.ac.uk/pub/databases/Pfam/Tools/) and eggnog mapper ^33^.

### Phylogenetic tree and estimation of divergence time

Protein datasets from eight representative salmon species were download from NCBI. Orthogroups (OGs) were identified using OrthoFinder ^73^. A total of 1,483 single copy orthologs were used to construct a phylogenetic tree using IQtree ^74^ with the model JTT+F+I+R7. Gene family expansions and losses were inferred using CAFE5 ^75^. The divergence time of each tree node was inferred using MCMCtree of the PAML package ^76^. The divergence time estimates were based on a known fossil record that gave an estimated divergence date of 6 MYA between *O. nerka* + *O. gorbuscha* and *O. keta* and 15-20 MYA as the divergence time between *Salmo*-*Oncorhynchus* genera ^36^.

### Whole-genome mapping and genotype calling

Whole-genome sequencing was conducted on two Amago, three Nagano Masu, and six additional *O. formosamous* individuals (Illumina platform, System). Raw paired-end reads were subjected to quality trimming using fastp (ver. 0.24.0) ^77^ with the following parameters “-q=30, -l=140, -5 -M=30 –cut_front_window_size=1, -r -M=30 –cut_right_window_size=4”. Subsequently, the processed reads were mapped to the reference genome assembly using BWA v.2.2.1 ^78^. Reads that were duplicated in the BAM files would be tagged with ‘DT’ using Picard MarkDuplicates of GATK v4.4.0.0 ^79^.

Variant calling of each salmon individual was performed using bcftools v1.19 ^80^ mpileup and call commands for minimum mapping quality of 30. Both monomorphic and polymorphic sites with read coverage were preserved in the initial output and subsequently merged to create one VCF file containing all individuals and all sites. The identification of biallelic single nucleotide polymorphisms (SNPs) was achieved through the use of ‘-m 2 -M2 -v snps’ options, followed by filters based on the criteria of ‘AC≥2 & F_missing ≤0.1’. To account for the variation in individual sequencing depth between studies, loci with sequencing depth less than 2 in more than 30% of the samples were excluded. Tentative false positive heterozygous SNPs exhibiting depth exceeding 175% of the individual genomic median were masked as missing genotype. The final dataset comprised 31.6 million SNPs after the above filtration process.

### Population genomic analysis

Principal component analysis was performed on the SNPs from the 25 largest scaffolds through PLINK v2.00a6LM ^81^. Population structure was analyzed using ADMIXTURE v1.3.0 ^82^ with fivefold cross-validation of K values from 2 to 8. According to the topology of the species tree, two *Oncorhynchus kisutch* individuals (SAMN36289694, SAMN36289695)^83^ were incorporated to the phylogeny analysis as outgroups to the *O. masou* species complex. The mapping rates of *O. kisutch* samples to the *O. formosanus* genome assembled in this study are higher than 98%. The best-fit nucleotide evolution model, TVMe, was determined by IQTREE v2.2.6 ^74,84^ based on the loci of filtered SNPs. A maximum-likelihood phylogenetic tree was then inferred on a 1000 ultrafast bootstrap approximation. The effects of SNP variants on protein-coding gene sequences were further annotated and classified as sites of loss of function (LoF) (gain or loss of stop codon and loss of start codon), missense, and synonymous (synonymous variant, start retained, stop retained variant) variants using SnpEff (ver. 5.2)^85^.

The demographic history was estimated using PSMC^86^. An additional three Illumina-sequenced individuals from Christensen *et al* ^28^ were mapped to genome assembly using bwa-mem (ver. 2.2.1)^87^ and the aligned mapped reads were converted to consensus sequence in FASTQ file using bcftools pipeline. The minimum (d) and maximum (D) coverage for vcfutils were set to 2 and 1.75X genome reads coverage. All of the parameters used for the PSMC program were at default with the exception of -𝜇 3.0e-09^88^ and -g 1^89^ .

### ddRADseq libraries preparation and data processing

Double-digest restriction-site-associated DNA sequencing (ddRADseq) was used to investigate the evolutionary history of Japanese and Taiwanese masou salmon populations, as well as the genetic diversity and demographic history of the Formosan salmon in Taiwan. Library preparation followed established protocols ^90,91^. Briefly, genomic DNA from each sample was digested using the restriction enzymes EcoRI and MseI (New England Biolabs, Massachusetts, USA), with ∼200 ng of DNA per individual. Adaptors containing unique barcode sequences were then ligated to the digested DNA fragments using T4 ligase (New England Biolabs). The barcoded DNA samples were individually purified using 1.8× AMPure XP magnetic beads (Beckman Coulter). PCR with eight cycles was performed on the purified fragments to incorporate Illumina sequencing adaptors. The resulting PCR products were pooled based on DNA concentration and underwent an additional round of magnetic bead-based size selection and purification. DNA fragments ranging from 300 to 500 base pairs (bp) were selected using 0.5× and 0.7× AMPure XP magnetic beads. In total, 83 samples were collected, including five Japanese samples (including two individuals of *O*. *m*. *ishikawae* (Amago) and three individuals of *O. m. masou* (Nagano masu) and 78 Taiwanese samples. The Taiwanese samples were distributed as follows: 21 from the Qijiawan creek, 17 from the Luoyewei creek, and 40 from the Hehuan creek. These samples were prepared into two sequencing libraries. Single-end sequencing with a read length of 150 bp was performed on an Illumina NovaSeq X platform at Tri-I Biotech, Inc. (New Taipei City, Taiwan).

The sequencing data were processed using the STACKS v2.62 pipeline^92,93^. First, the process_radtags program in STACKS was used to demultiplex the sequencing data with default settings. Reads were retained if they contained an intact EcoRI RAD site, had a unique barcode, and did not include Illumina adapter sequences. The trimmed reads were then mapped to the *O*. *formosanus* reference genome using the reference-mapping process in STACKS. Specifically, the reference genome was indexed using the bwa index command^94^, and each sample was aligned using the bwa mem mode. The mapped results were then compressed while preserving the headers into individual BAM files using Samtools^80^.

Following the read alignment, two datasets were generated for downstream analysis. The first dataset included both Japanese and Taiwanese samples and was used to examine the evolutionary history of Japanese and Taiwanese masou salmon. The second dataset contained only Taiwanese samples and was used to estimate the genetic diversity and demographic history of the Formosan salmon. Loci were assembled using the ref_map.pl wrapper script in STACKS, and the populations module was used to export the results. To ensure data quality, loci were retained only if they were present in at least 80% of individuals per population, shared across all populations, and had a minor allele count of at least three.

### Population tree estimation and species delimitation

We used the Bayesian Phylogenetics and Phylogeography program (BPP, version 4.7.0)^37,95^ to estimate a species tree at the population/taxon level, including divergence times, and to delimit evolutionarily independent entities under the multi-species coalescent model^38,96^. Given that our sampling design did not include outgroup samples, we first used the BPP program to estimate the species tree topology while simultaneously delimiting evolutionarily independent entities using the A11 model. Next, we fixed the best estimated species tree topology and used it to estimate genetic diversity (theta) for each terminal taxon, as well as the ancestral node, and coalescent times (tau) for each internal node, applying the A00 model.

A time-calibrated diversification history was then generated, using a generation time of 1 year and a spontaneous genomic mutation rate of 1.06 x 10^-8^ bp/generation^97^, via the bppr package (https://github.com/dosreislab/bppr) and the results from the A00 model of BPP analyses. Furthermore, the estimated theta and tau values were subsequently used to calculate the genealogical divergence index (GDI)^98^ for different divergence events on the species tree. The GDI values have been shown to capture nuances that represent various stages of the speciation continuum, providing a more evolutionarily informed framework for species delimitation^98,99^. A practical application of the GDI was recently carried out in a large-scale study on darter fishes in the U.S. for conservation purposes^40^.

To facilitate the tractability of computational power while ensuring the repeatability and reproducibility of our results from BPP analyses, we generated three independent datasets for each BPP analysis, with each dataset containing 3000 randomly selected RAD loci from across the genome. We consistently included the three Amago and two Nagano Masu samples. For the Formosan salmon populations, we randomly included three individuals from the Qijiawan population and three individuals from the Hehuan population, for a total of six individuals. To generate control files for the BPP analyses, we used an online tool provided by the authors of the BPP program (https://brannala.github.io/bpps/#/). This online tool also helped with specifying appropriate prior settings for the theta and tau parameters based on the empirical sequence data. The BPP analyses were run for 5 × 10^5^ generations, with samples taken every 5 generations, and a burn-in period of 1 × 10^4^ generations was applied. The convergence of the analyses was assessed by ensuring that the effective sample sizes (ESSs) for all estimated parameters were greater than 200.

### Population genetic structure of the Formosan salmon

To investigate the genetic relationships among Formosan salmon populations, we conducted a principal component analysis (PCA). First, we imported VCF files into R using the read.vcfR function from the vcfR package^100^. To mitigate the effects of linkage among SNPs, we filtered SNPs using the distance_thin function in the SNPfiltR package^101^, setting a minimum distance of 1 Mb (min.distance = 1,000,000). Subsequently, the filtered SNP data were converted into genind objects using the vcfR2genind function, with populations defined based on eight collection sites across three streams. To handle missing data, we applied the scaleGen function from the adegenet package^102^, replacing missing values with the mean (NA.method = "mean"). Finally, we performed PCA on the transformed genind dataset (center = TRUE, scale = TRUE) using the dudi.pca function from the ade4 package^103^.

### Estimating the genetic diversity and level of inbreeding in the Formosan salmon

To calculate the genetic diversity of all the individuals across the three stream populations in Taiwan, we assessed the multi-locus heterozygosity (MLH) ^104^ using the *MLH* function within the *inbreedR* package^105^ in R. The estimated MLH values were then visually represented using bar charts, with each bar representing the heterozygosity of each individual.

To estimate runs of homozygosity (ROHs) for each individual, we used PLINK v1.9^106^ to calculate the number and length of ROHs (>1000 Kb) across the genome. ROHs were estimated using two different window sizes, 5 and 10 SNPs (-- homozyg-window-snp 5; --homozyg-window-snp 10). The parameters for ROH estimation were adapted from previous studies^107,108^ as follows: allowing one heterozygous SNP per window (--homozyg-window-het 1), permitting one missing call per window (--homozyg-window-missing 1), and setting the percentage of overlapping windows required to classify a SNP as part of a homozygous segment to 1% (--homozyg-window-threshold 0.01). Finally, segments were considered homozygous if they exceeded 1000 Kb in length (--homozyg-kb 1000).

### Estimating current effective population sizes and reconstructing the demographic history

To estimate the current effective population size of the three geographical taxa, we applied NeEstimator V2^109^ using the *gl.LDNe* function^110^ in dartR package in R^111,112^. The genlight objects for downstream analyses were imported from the VCF files using the *gl.read.vcf* function in dartR, with populations defined by the three streams. To account for the linkage among SNPs from the same chromosome, we specified the argument *pairing* in the *gl.LDNe*, which limited the use of paired loci only when they were from different chromosomes when estimating the effective population size. We estimated the effective population size both with and without empirical corrections by using the *naïve* argument in the *gl.LDNe*. The effective population size for each stream populations was estimated under the assumption of random mating within each population, while considering only alleles with a frequency higher than 0. Specifically, we do not observe genetic structure within populations, and the populations appear to be relicts of hypothesized recent local extirpation events; therefore, assuming a panmictic population, or random mating system, is biologically appropriate.

To reconstruct the recent demographic history of Formosan salmon from the Qijiawan and Hehuan Streams, we applied a linkage disequilibrium-based method implemented in GONE^113^. The population from the Luoyewei Stream was excluded due to its high level of inbreeding, as excessively long ROHs would lead to erroneous estimates in the software. The sequence data were treated as unknown phase, and the estimation window was set to 100 generations, divided into 100 time bins, with each bin representing one generation. The maximum recombination rate between analyzed SNP pairs was set to 0.05 (hc = 0.05), and a maximum of 50,000 SNPs per chromosome was analyzed. For the GONE analyses, we used a generation time of 1 year and the average recombination rate of salmon (1.09 cM/Mb)^114^. Additionally, to assess run-to-run variability in GONE results, we conducted 100 independent runs using the average recombination rate of salmon.

### Non-lethal field sampling and age estimation

Given the endangered status of Formosan landlocked salmon (*Oncorhynchus formosanus*) and strict legal restrictions on lethal sampling, we used low- and non-invasive monitoring techniques from 2021 to 2024. Between 2021 and 2023, individuals were sampled using hook-and-line fishing; however, this method exhibited size selectivity that could bias estimates of population structure. To address this limitation, in 2024 we switched to snorkel-based underwater visual surveys with scale-referenced imaging to non-invasively estimate body length distributions. During the late breeding season (November-December 2024), individuals were collected for morphometric and age data using non-lethal methods: night snorkeling capture in Chichiawan Creek (QIJ) and electrofishing in Hehuan Creek (HEH). Sampling periods, methods, and summaries of metrics are detailed in Table S12 and Table S13.

For masu salmons, scales have been shown to provide age estimates that are not only consistent, but in many cases more accurate than those derived from otoliths^115,116^. Due to legal prohibitions on lethal sampling for otolith extraction (Taichung City Agricultural Bureau 2021), we used scales as the primary material for age inference. Previous validation experiments report scale-based age accuracy between 90-96% using standard sampling protocols^116^. We collected scales from the preferred body region (posterior to the dorsal fin and anterior to the anal fin, above the lateral line, in the 2nd-3rd scale rows^115,117^). After rinsing and air drying, five good quality scales per individual were selected. These were cleaned and pressed using a roller device to produce acetate impressions. The rings were examined by reflected light microscopy at 30-40x. Annual rings were identified based on abrupt transitions from widely spaced summer rings to densely packed winter rings. False rings and spawning checks were excluded from age counts according^115,116^. A single trained reader performed all scale readings to ensure internal consistency.

Observed lengths from underwater surveys were corrected for light refraction by dividing raw values by the refractive index of water (1.33). To further correct for detection bias across size classes, paired snorkel and electrofishing surveys were conducted at HEH in 2024. Electrofishing, assumed to be less biased across size classes, provided a reference distribution. Length-specific detection ratios were calculated by comparing the frequency distributions of the two methods in overlapping size bins (e.g., 37.6-75.2 mm, 75.2-112.8 mm). Ratios < 1 indicated underestimation by snorkeling, and snorkel-based length counts were adjusted accordingly. Corresponding detection distributions and correction curves are shown in Figures S15 and S16.

Snorkel observations provided the number of fish within specific length intervals (Fig. S17), but did not provide individual-level measurements. To reconstruct individual body lengths, we implemented a probabilistic sampling approach. For each observation of N individuals within a length interval [Lmin, Lmax], N measurements were randomly sampled with replacement from the corresponding donor pool of scale-measured individuals. Sampling was constrained by stream and sampling period (e.g., only samples from HEH_11-12_ were used to reconstruct HEH November-December data). When sample sizes in a donor pool were insufficient, sampling was extended incrementally to similar intervals. This resulted in a reconstructed distribution of individual lengths for each snorkel dataset.

### Estimation of demographic parameters

Because age-specific capture probability (𝑞_age_) cannot be measured directly, we adopted a pragmatic two–gear strategy in which backpack electrofishing served as the least age–biased reference method. Independent field studies on salmonids demonstrate that electrofishing efficiency rises with body size but plateaus at a high, nearly constant level once fish pass the early–juvenile stage^118,119^, rendering capture probability effectively age–invariant for fully recruited age classes. We therefore used electrofishing counts to approximate an equal–catchability age distribution and, for the same stream reaches, calculated length–specific detection ratios between electrofishing and snorkel surveys. These ratios were applied as correction factors to snorkel counts, compensating for the under–representation of the smallest size classes. After correction, residual differences in 𝑞_age_ among fully recruited ages were negligible, satisfying the Chapman–Robson assumption of equal catchability.

To assign ages to individual fish, we implemented a two–step procedure that propagates uncertainty in the length–age relationship. First, 1,000 bootstrap resamples were drawn from the empirical length–age data set to obtain a distribution of the mean (𝜇_𝑖_) and standard deviation (𝜎_𝑖_) for each age class (Fig. S18). For Qijiawan (QIJ), which lacked prior age data, we used maximum likelihood: for each fish of length 𝑙 we evaluated dnorm(𝑙; 𝜇_𝑖_, 𝜎_𝑖_) over all ages and assigned the age with the highest likelihood. For Hehuan (HEH), where electrofishing supplied reliable priors, we adopted a Bayesian approach in which posterior probabilities were proportional to dnorm(𝑙; 𝜇_𝑖_, 𝜎_𝑖_) 𝑃(Age = 𝑖). Repeating this across the 1,000 resamples yielded age–class proportions and 95 % confidence intervals for every site–month combination (Table S9).

Three demographic quantities were then estimated: (i) annual recruitment of age– 0+ fish, (ii) seasonal survival rates (January→November and November→June), and (iii) age–specific fecundity coefficients. All site-level quantities were first calculated on an area-specific basis (ind. 1000m⁻²) and then aggregated to the reach or stream scale as area-weighted means. Seasonal, age-specific survival was estimated by tracking the same cohort between successive surveys (matching age i at the first survey to age i + 1 at the next), rather than by cross-sectional comparisons among ages. We analysed two intervals: January→November (10 months; typhoon season) and November→June (7 months; baseline). Site-level estimates were aggregated to the stream scale using area-weighted means, then standardized to monthly and annual scales to make intervals comparable. Because survival can vary with age, all estimates were computed and reported by age class. Estimates were flagged as unreliable when extremely low or high (below 0.05 or above 0.95) or when the bootstrap 95% CI was very wide (width > 0.60). Flagged cells were replaced with values sampled from the empirical distribution of age-2 survival in the paired stream during the same season, preserving seasonal and age structure while avoiding unstable ratios. Replicate-level results were summarized as stream means with 95% confidence intervals..

To contrast population structure before and after the 2024 typhoon sequence, we reconstructed age vectors for January 2024 (baseline) and January 2025 (post– typhoon) by temporal back–tracking. The baseline series was anchored to the last comprehensive survey in June 2024, preceding the typhoon season. A monthly baseline survival rate (𝑆_base_) was extracted from mark–recapture data collected between November 2023 and June 2024, and the June abundance vector was propagated backwards five months,

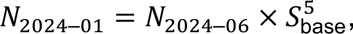

The post–typhoon series used the November–December 2024 survey as its anchor; a winter survival rate (𝑆_winter_) for the November–January interval was estimated and applied for one–month back–projection,

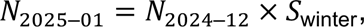

Counts of fish aged ≥ 3 y—already near the species’ maximum longevity—were left unadjusted at both anchor dates.

Recruitment (R) at each anchor month was obtained from a Chapman–Robson catch-curve analysis applied to the reconstructed age distributions. Instead of using the method to estimate mortality, we interpreted the log-linear catch curve intercept (back-extrapolated to age 0) as an index of initial cohort abundance, i.e. relative recruitment strength ^120^. This approach relies on the standard catch-curve identity in which the intercept reflects q ⋅ N0 under the assumptions of equal catchability and approximately constant mortality across fully recruited ages. Catchability was equalized by using electrofishing as a reference gear and applying length-specific detection corrections to snorkel counts (see above). Because disturbance years can violate strict steady-state assumptions, we explicitly treat R as a relative index rather than an absolute number, using it only for between-stream and between-scenario comparisons. Replicate-level estimates were aggregated to stream-level means and 95% confidence intervals after area-weighting sites.

We then linked recruitment indices to mature-female density through a scaling factor B. Specifically,

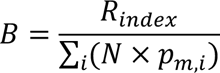

where 𝑁_𝑖_is the abundance of age class 𝑖, and 𝑝_𝑚,𝑖_ is the empirically derived proportion of mature females at age 𝑖. Thus, 𝐵 represents the number of recruits per mature-female equivalent. Age-specific fecundity coefficients were then defined as 𝐹_𝑖_ = 𝑝_𝑚,_ 𝐵, representing the expected contribution of age-𝑖 females to the recruitment index. For QIJ in 2024, a correction factor of 0.719 was applied to account for spatial sampling bias associated with an upstream, high–productivity reach^23^. Fecundity parameters were estimated separately for January 2024 (pre-typhoon) and January 2025 (post-typhoon back-track). Importantly, these are index-scale fecundities rather than absolute egg counts, consistent with age-structured modelling that assigns age-specific reproductive weights to drive population projections ^121^. This framework provides the demographic inputs for the stochastic simulations described below.

### Stochastic simulation of population dynamics

To assess the long-term effects of typhoon disturbance on salmonid population viability, we constructed a discrete-time, age-structured simulation model using Leslie matrices. In our simulations, we defined a ’typhoon year’ as any year in which at least one typhoon event caused observable impacts on stream habitat and salmon populations during the July-October period, based on historical records from the Central Weather Bureau. We treated typhoon occurrence as a binary variable (present/absent) rather than incorporating intensity gradients, as our demographic data did not allow for parsing intensity-specific effects. The model incorporated two distinct demographic scenarios representing typhoon and non-typhoon years, with parameters derived from empirical field estimates. The age structure included four classes (0+-3+), with age-specific fecundity coefficients and survival rates parameterized from field data. Fecundity vectors were scenario-dependent: "noTyphoon" scenario used parameters derived from January mature females contributing to July recruitment, while "afterTyphoon" scenario used parameters from post-typhoon population reconstruction. Annual survival rates were calculated from two distinct seasonal periods: baseline survival rates were derived from November→June transitions (7-month period representing non-typhoon conditions), while typhoon-impacted survival rates were estimated from January→November transitions (10-month period encompassing the typhoon season). These period-specific survival estimates were then standardized to annual rates for incorporation into the Leslie matrix framework.Carrying capacity was set at 1,500 individuals per 1,000 m², with proportional scaling applied when simulated populations exceeded this threshold. The simulation proceeded as follows: (1) determination of typhoon occurrence for the focal year; (2) selection of appropriate survival and fecundity parameter sets; (3) construction and application of the corresponding Leslie matrix; (4) density regulation if population exceeded carrying capacity; and (5) transition to the next annual cycle. Initial population structure was derived from November 2024 field observations for both Qijiawan Creek (QIJ) and Hehuan Creek (HEH). Simulations were replicated 1,000 times for each typhoon probability scenario over 20-year periods, generating age-structure trajectories and total population size distributions for comparative analysis.

For every replicate we extracted the dominant eigenvalue (λ) of the corresponding Leslie matrix and summarised its distribution by the mean and 95 % CI to compare ‘no-Typhoon’ and ‘after-Typhoon’ scenarios. Quasi-extinction was defined as a total population density ≤ 10 indvidual per 1,000 m². Extinction counts across 1,000 replicates were compared between streams by Fisher’s exact test, yielding odds ratios (OR) with exact 95 % CI. To test for a stream × typhoon-probability interaction, we fitted a logistic GLM to the binary extinction outcome, omitting p = 0.9 to avoid complete separation.

### Research permits and ethical compliance

All fieldwork conducted between 2021 and 2024 was approved by relevant regulatory agencies, including the Taiwan Forestry and Nature Conservation Agency (permits: 1132401290, 1130218297, 111701591), Shei-Pa National Park (1110002834, 1130003015), and Taroko National Park (1110003283, 1131005811). All procedures complied with the Wildlife Conservation Act and National Park regulations. Capture methods included non-lethal snorkel surveys, hand netting, fly fishing, and limited electrofishing. All individuals were anesthetized with 2-phenoxyethanol (0.1-0.3 mL L-¹), handled within 3 minutes, and released after full recovery. Only two individuals were sacrificed for genomic sequencing under the approval of Taiwan Forestry and Nature Conservation Agency, Shei-Pa National Park and Academia Sinica IACUC (24-09-2284). All research followed the principles of replacement, reduction and refinement (3Rs).

## Data Availability

All sequences generated from this study were deposited on NCBI under BioProject PRJNA1277495 and accession numbers of individual samples can be found in Table S1.

## Acknowledgments

We thank Dr. Chyng-Shyan Tzeng (National Chung Hsing University, Taiwan), Dr. Shih-Pin Huang (Biodiversity Research Museum, Academia Sinica, Taiwan) and Yu-Ching Liu (Biodiversity Research Center, Academia Sinica, Taiwan) for their assistance.

## Supplementary Figures

**Fig. S1.**
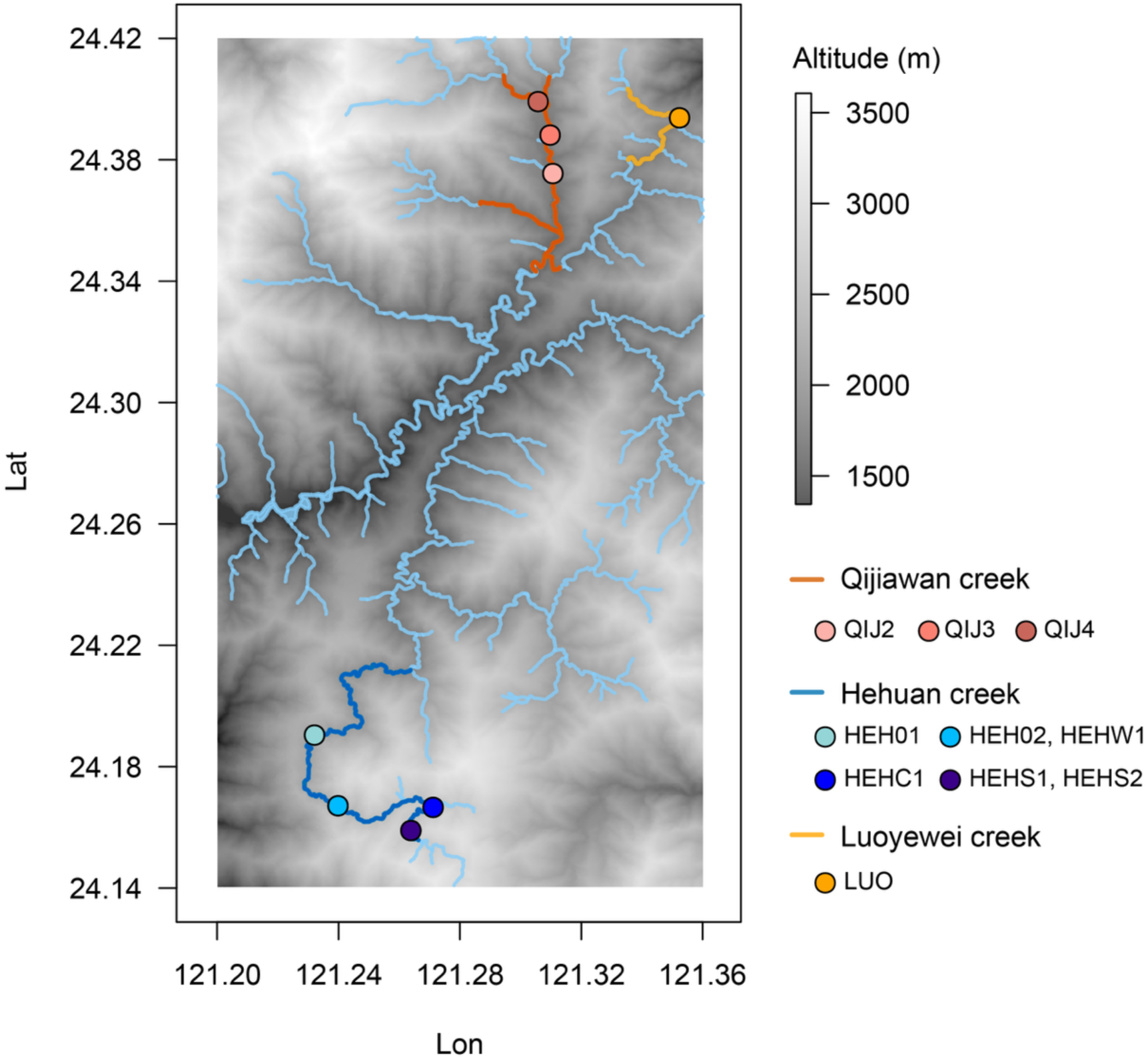
Geographic distribution and sampling locations of *Oncorhynchus formosanus* populations in Taiwan’s Central Mountain Range. The map shows the three creek systems harboring extant O. formosanus populations at elevations between 1,500-3,500 m. Qijiawan Creek (orange line) in the northern portion contains the historically protected population with sampling sites QIJ2, QIJ3, and QIJ4. Hehuan Creek (blue lines) in the southern portion includes both upstream sites (HEH01, HEHS1, HEHS2; dark blue circles) and downstream sites (HEH02, HEHW1; light blue circles) where reintroduction occurred in 2017. Luoyewei Creek (yellow line) in the eastern portion represents the hatchery-supplemented population (LUO). The grayscale gradient indicates elevation, with darker shades representing lower elevations. All populations are located within the upper tributaries of the Dajia River basin in central Taiwan.

**Fig. S2.**
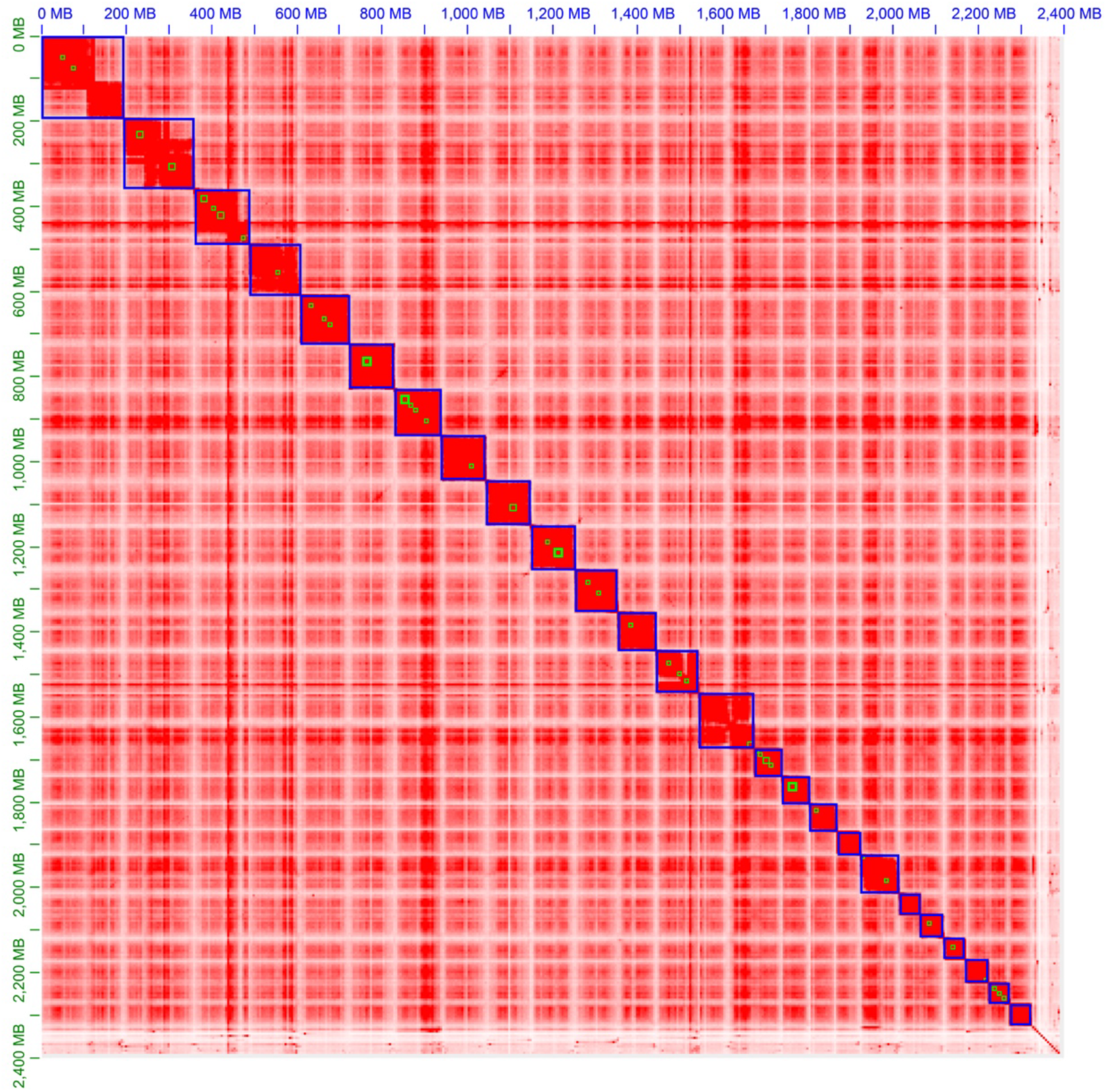
Genome-wide Hi-C contact map of the *O. formosanus* genome assembly. The Hi-C interaction heatmap illustrates the genome-wide three-dimensional organization and chromosomal-level scaffolding of the *O. formosanus* genome. Each blue box represents an individual chromosome-scale scaffold identified in this study. Stronger interaction signals are indicated in red, representing frequent physical interactions within chromosomes, while weaker or infrequent interactions appear in lighter colors.

**Fig. S3.**
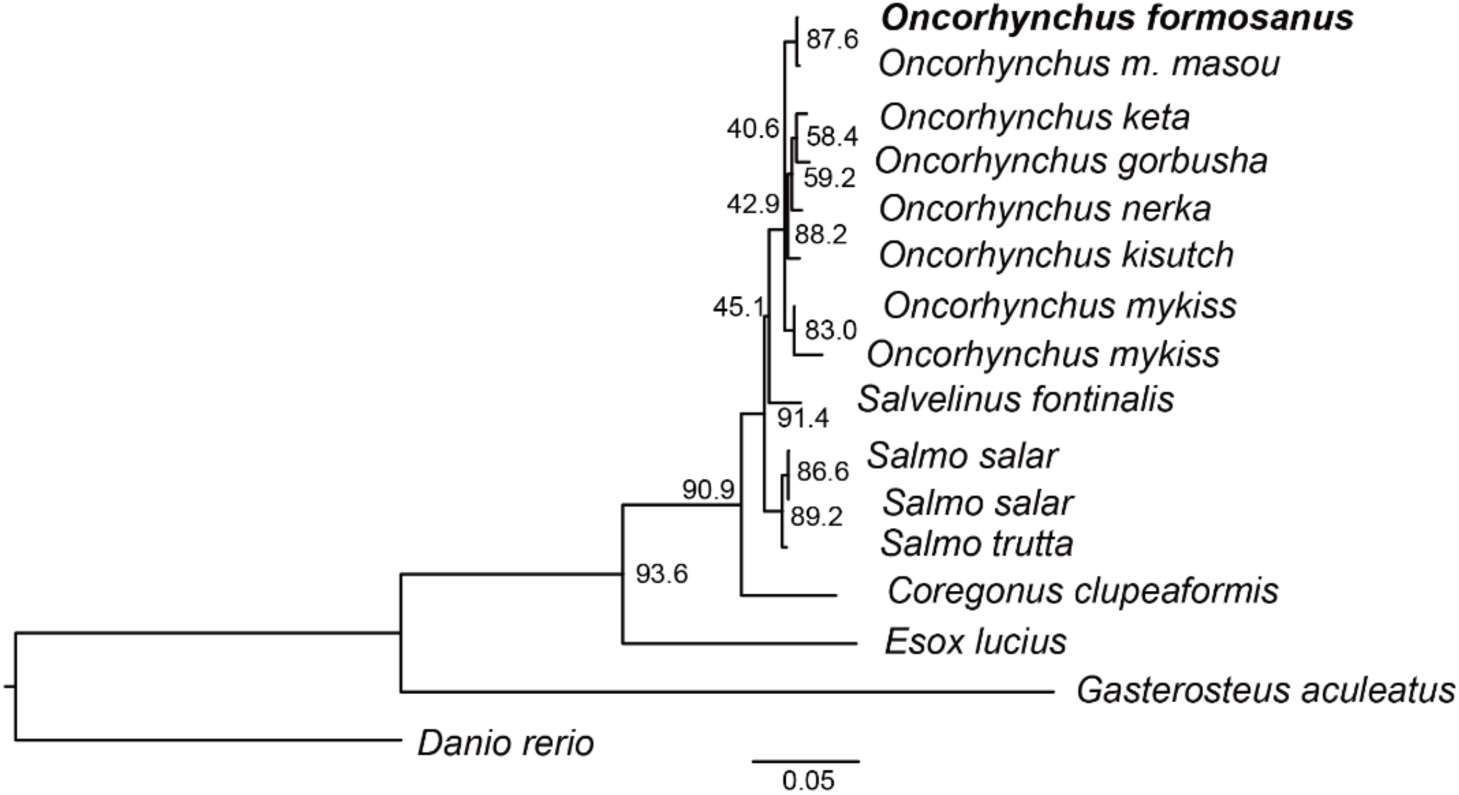
Coalescent-based species tree inferred by ASTRAL for Oncorhynchus species. Species phylogeny constructed using the coalescent-based approach implemented in ASTRAL from 1,483 single-copy orthologs. Branch support values represent local posterior probabilities, reflecting concordance among gene trees. This analysis provides the same topology and corroborates relationships among *Oncorhynchus* species compared to the species tree from concatenated alignments of Fig. 1b.

**Figure S4.**
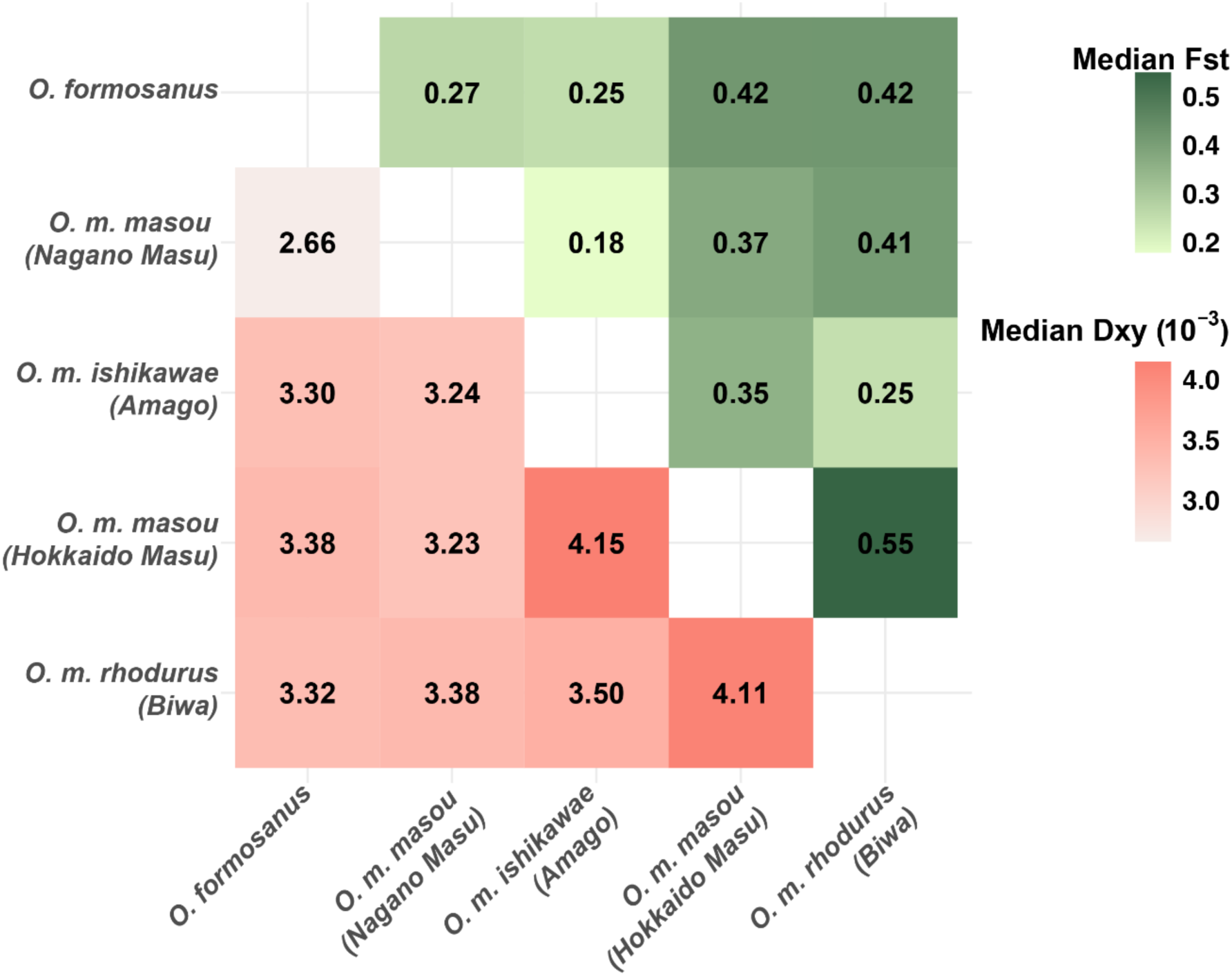
Pairwise genetic differentiation among masu salmon lineages. Heatmap illustrating pairwise genetic differentiation (Fst, green shades) and absolute nucleotide divergence (Dxy, red shades) among five *Oncorhynchus masou* lineages.

**Fig. S5.**
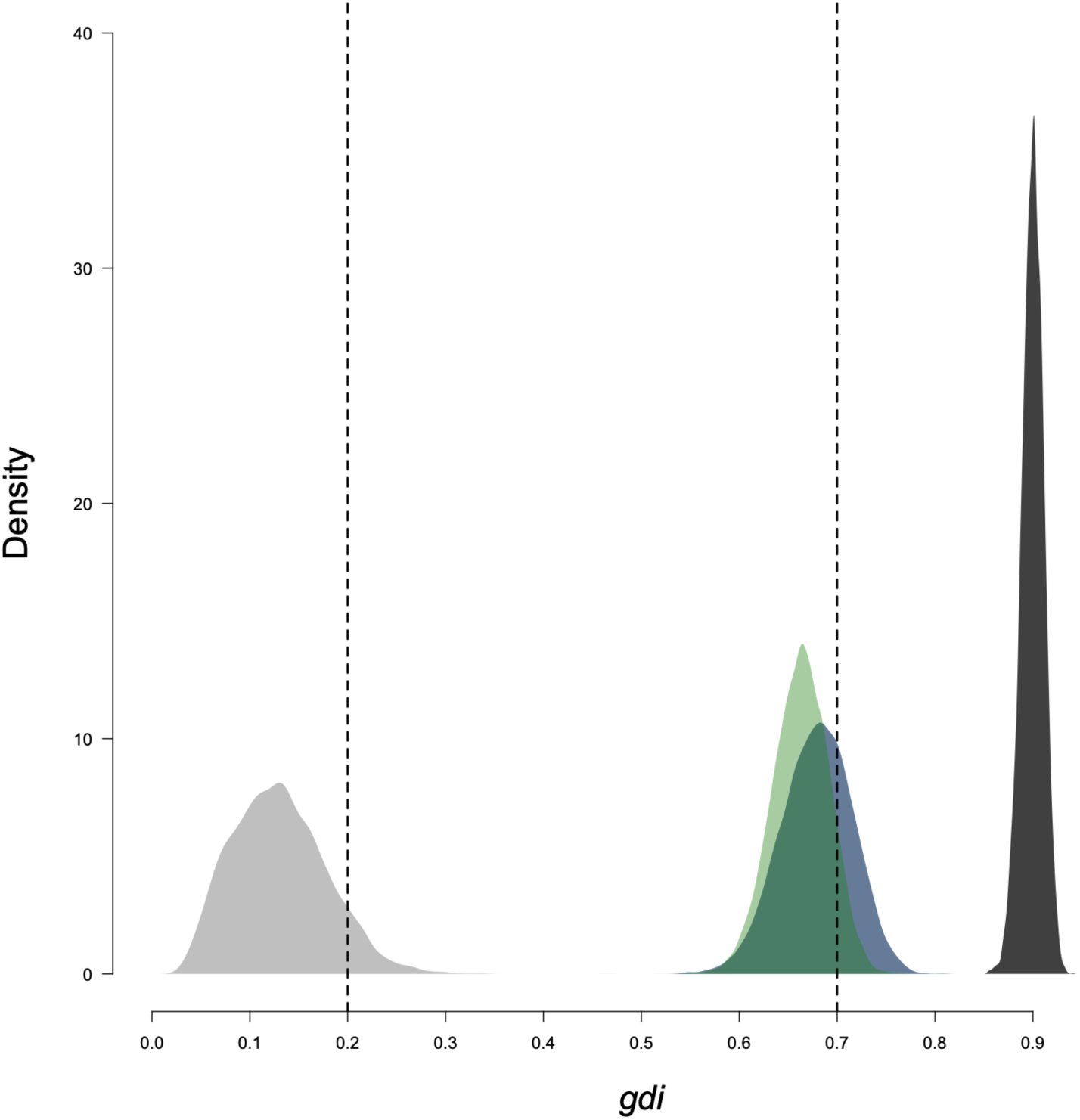
Genealogical Divergence Index (GDI) analysis of Taiwanese and Japanese salmon lineages. **The** genome-wide estimates of genealogical divergence index (GDI) reveals substantial genetic differentiation between Taiwanese (*O. formosanus*) and Japanese (*O. masou*) salmon lineages (the value was around 1.0; dark grey). The estimated GDI values between the two Japanese salmon lineages were highlighted in green and blue colors, where the values were around 0.7. High GDI values (>0.7) indicate strong genealogical divergence, implying long-term isolation and lineage differentiation. In contrast, the estimated GDI between the two Taiwanese population (highlighted in light grey) was smaller than 0.2, which suggests neglectable genetic divergence between the two populations.

**Fig. S6.**
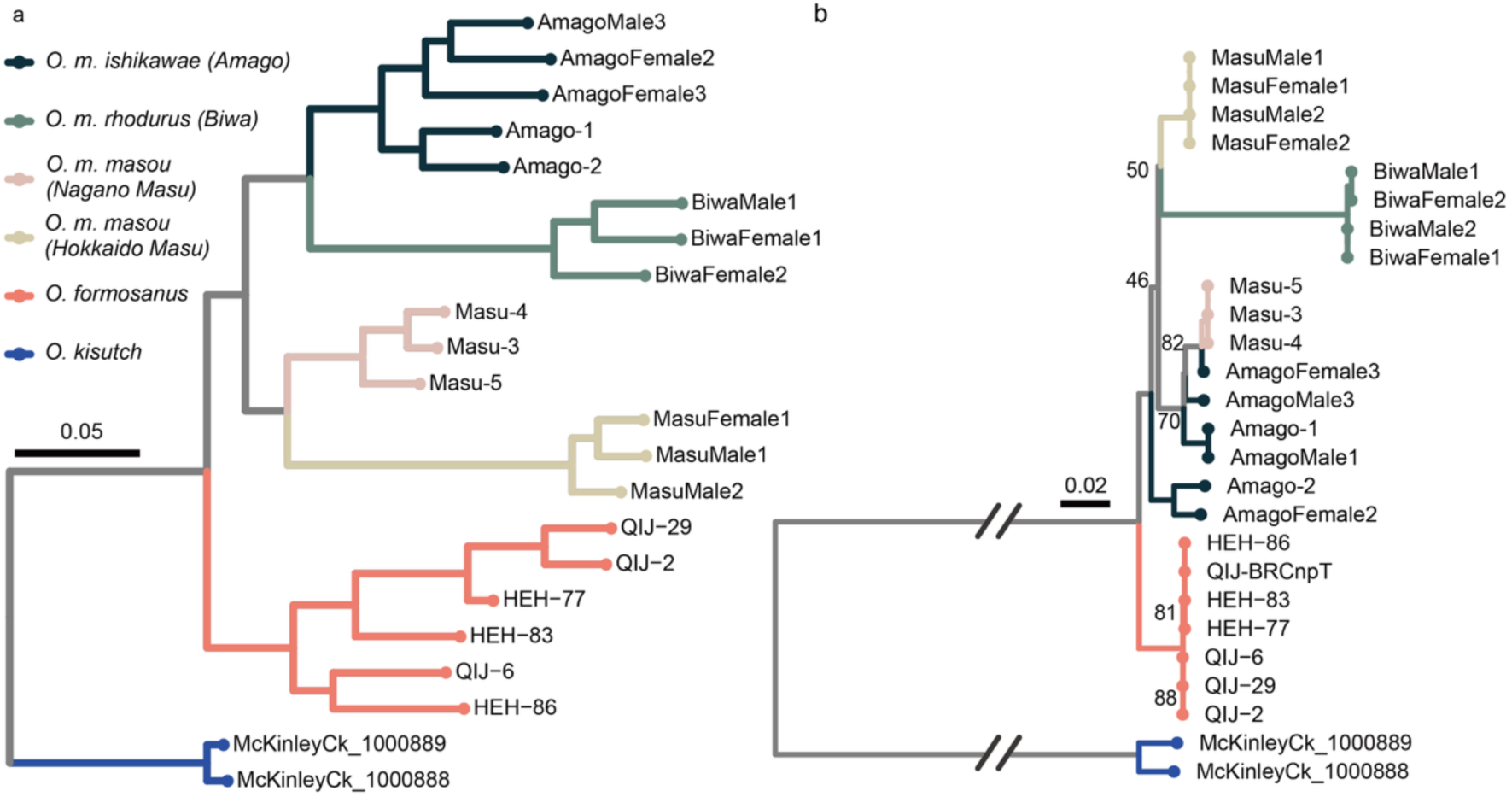
Population phylogeny of Taiwanese and Japanese salmon lineages inferred from nuclear genome-wide and mitochondrial SNPs. (a) Maximum likelihood phylogeny based on genome-wide SNP data from representative individuals of Taiwanese (*O. formosanus*: QIJ, HEH) and Japanese (Amago, Biwa, Nagono Masu and Hokkaido Masu) salmon populations, rooted with *O. kisutch* (coho salmon) as the outgroup. Branch lengths represent nucleotide substitutions per site. (b) Corresponding mitochondrial genome phylogeny inferred from mitogenome SNP data, also rooted with *O. kisutch*. Bootstrap support values (percent) for major branches are indicated. Together, these phylogenies illustrate consistent genetic differentiation patterns between nuclear and mitochondrial genomes across salmon lineages.

**Fig. S7.**
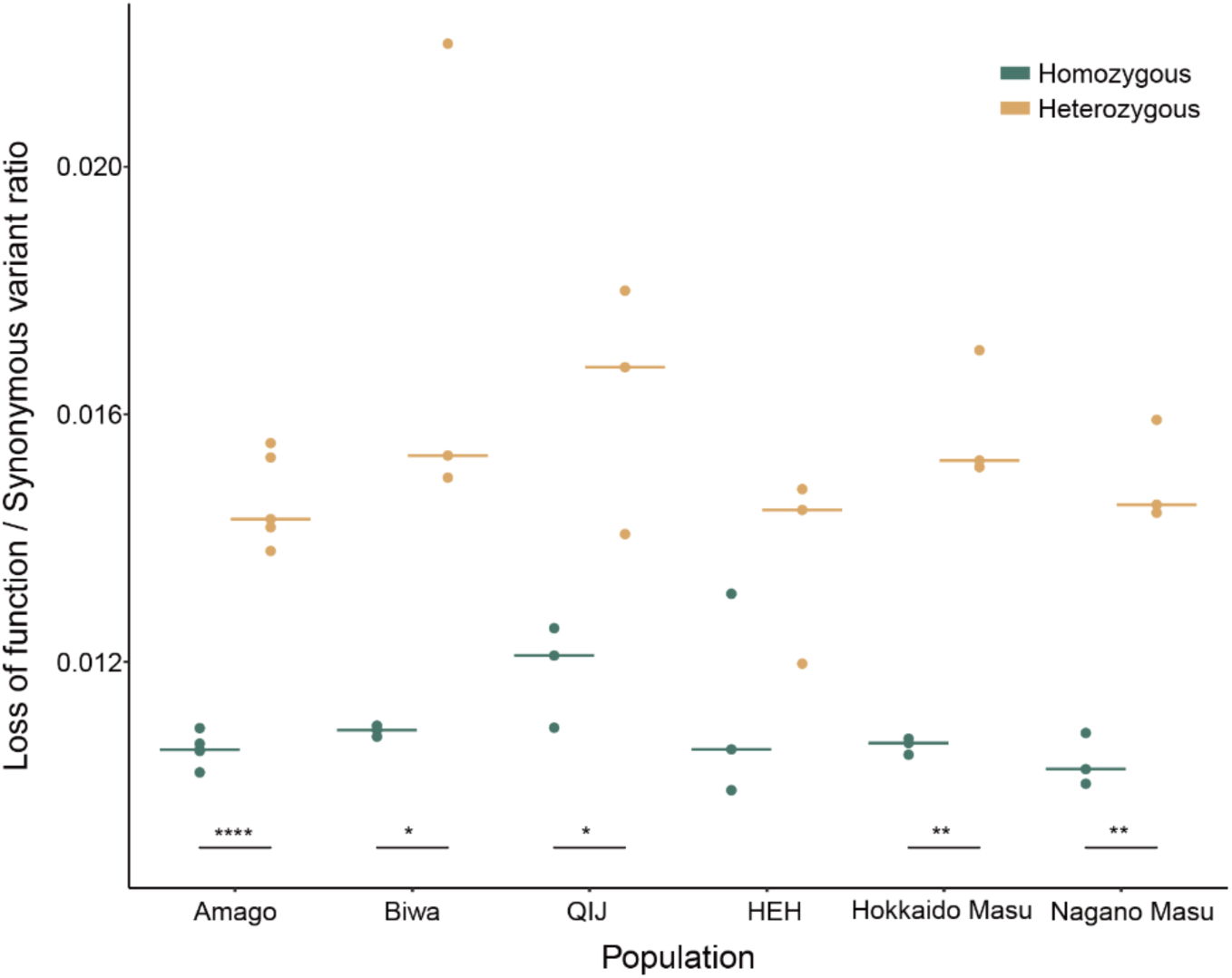
Relative proportions of loss-of-function to synonymous variants in homozygous and heterozygous states across salmon populations. Scatter plot showing the ratio of loss-of-function (LOF) variants to synonymous variants in both homozygous (green) and heterozygous (orange) states among salmon populations: Amago, Biwa, QIJ (Qijiawan), HEH (Hehuan), Hokkaido Masu, and Nagano Masu. Horizontal bars indicate mean values for each population. Heterozygous LOF variants consistently exhibit higher ratios compared to homozygous LOF variants across all populations, aligning with expectations of increased accumulation of deleterious alleles and elevated homozygosity in populations with smaller effective population sizes. Asterisks denote statistical significance of the difference between homozygous and heterozygous states (*P<0.05, **P<0.01, ****P<0.0001).

**Figure S8.**
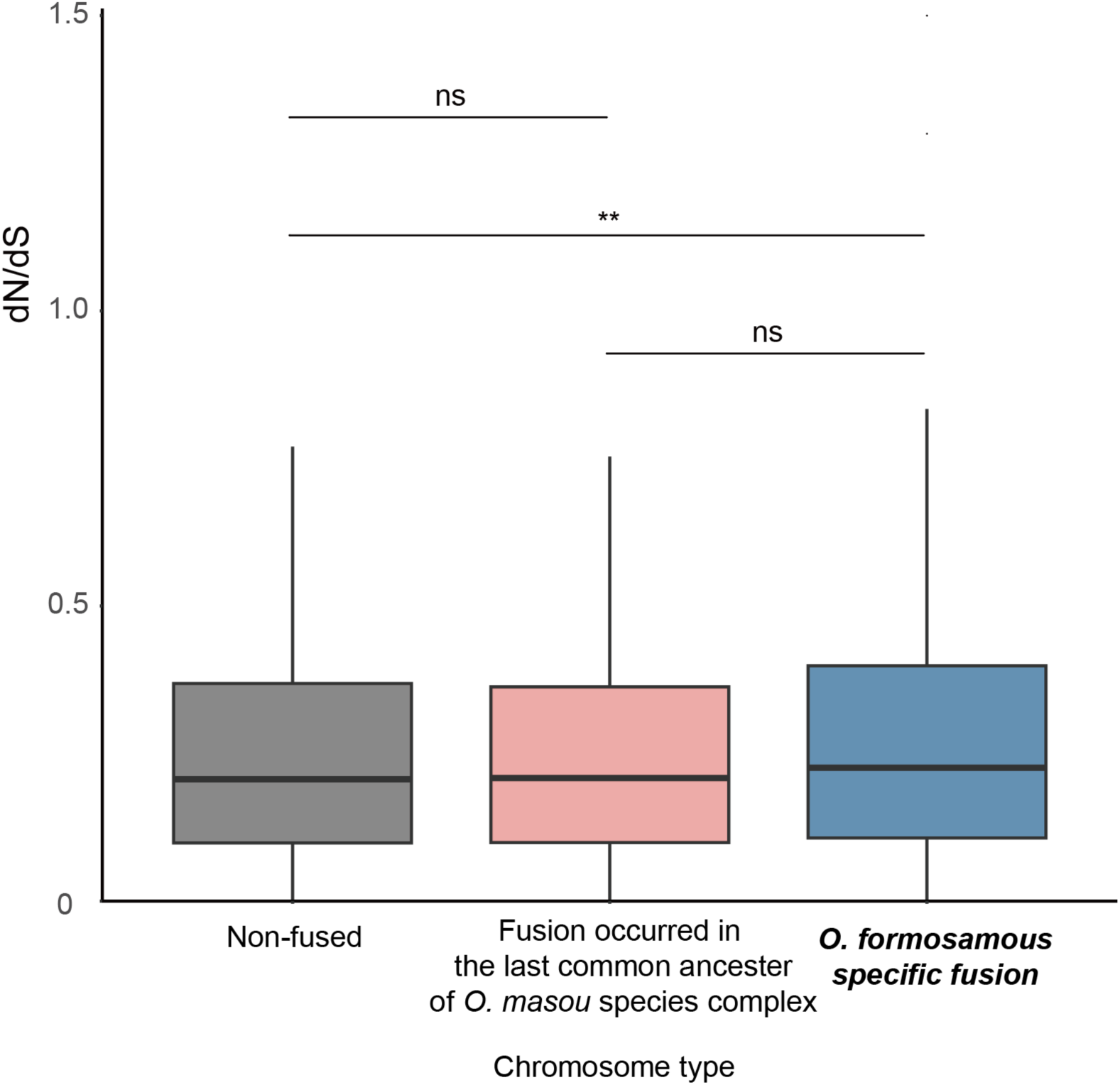
Comparison of evolutionary rates (dN/dS ratios) across different chromosome categories in *O. formosanus*. Boxplot showing the distribution of nonsynonymous-to-synonymous substitution rate ratios (dN/dS) for genes located on chromosomes classified as non-fused, fused in the last common ancestor of the *O. masou* species complex, or specifically fused in *O. formosanus*. "ns" denotes no significant difference, ** denote P < 0.01 in Wilcoxon rank sum test.

**Fig. S9.**
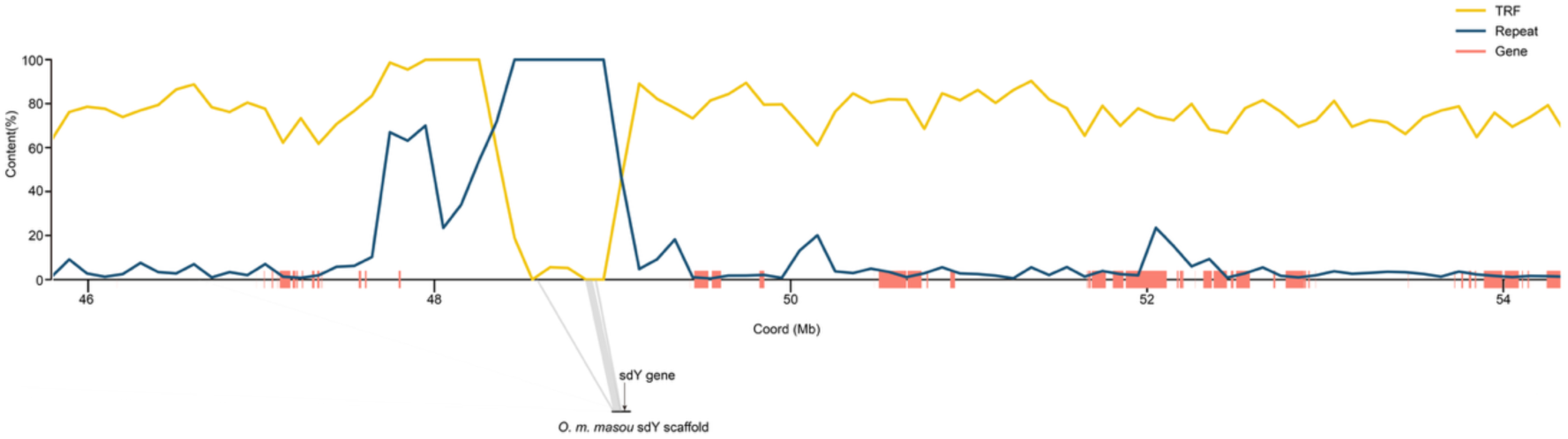
Alignment of the female-derived *O. m. masou* sdY-containing contig to chromosome 13. Dot plot visualization illustrating the unique alignment of the female-derived *O. m. masou* contig containing the sex-determining gene (*sdY*) to the central genomic region (∼50 Mb) of chromosome 13. Alignment identity is indicated by a color scale, with higher identity in red. This confirms the precise chromosomal location of the sex-linked region at an ancestral chromosome fusion site.

**Fig. S10.**
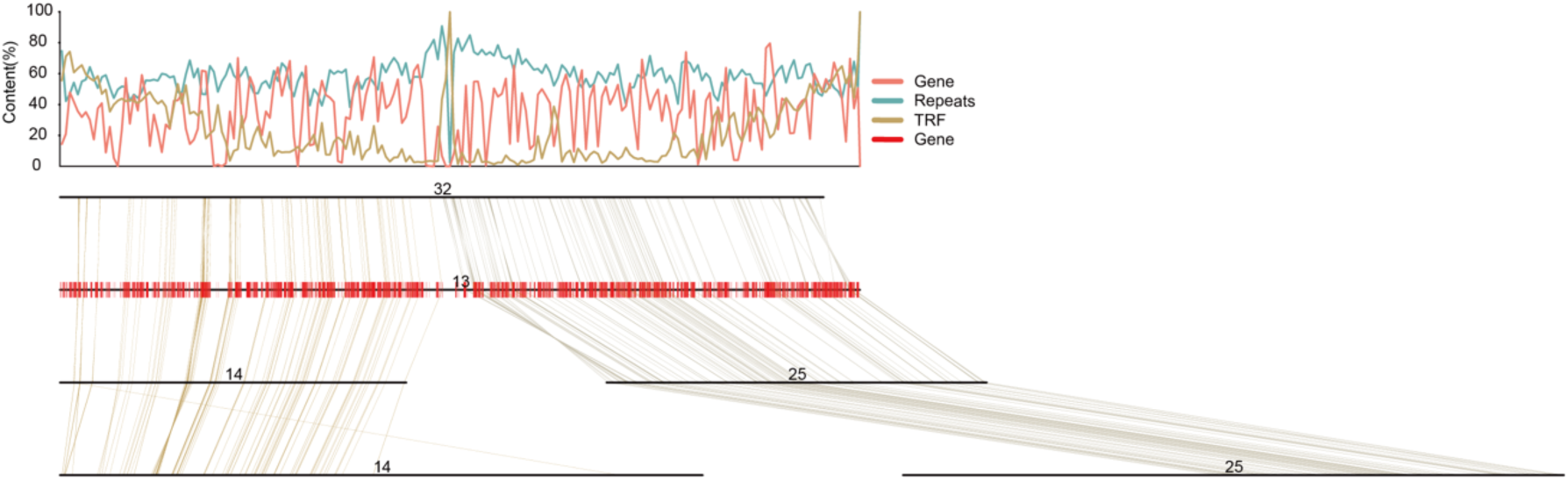
Comparative alignment showing ancestral chromosome fusion between chromosome 13 of *O. m. masou* and chromosomes 14 and 25 of *O. mykiss*. Visualization illustrating the genomic correspondence and alignment between chromosome 13 of *O. m. masou* and chromosomes 14 and 25 in rainbow trout (*O. mykiss*). The ancestral chromosome fusion site includes enriched regions of repetitive sequences, notably 281-mer satellite repeats, indicative of centromeric or neocentromeric activity and consistent with chromosome fusion events. Sequence similarity is depicted by color intensity.

**Fig. S11.**
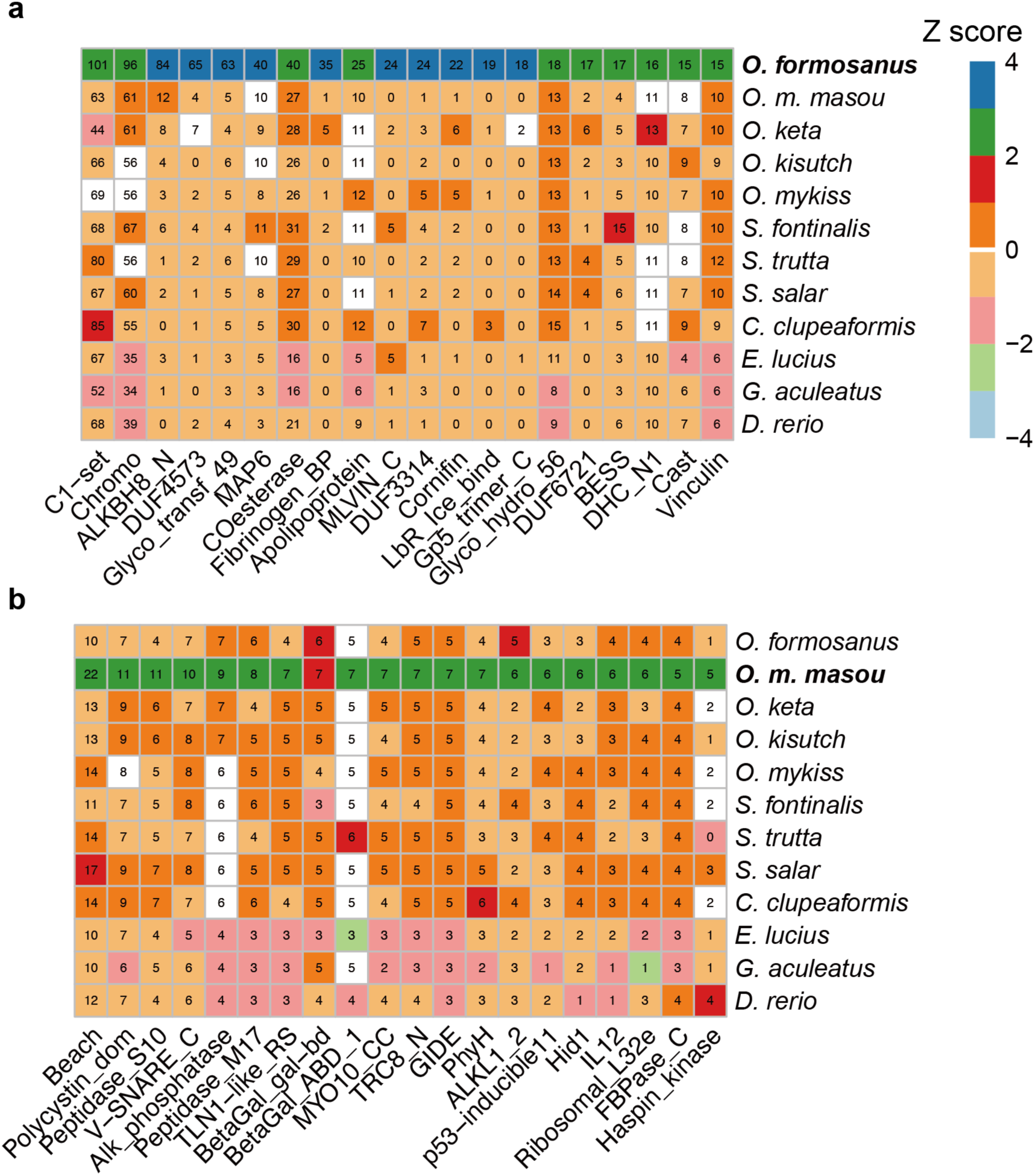
Protein family (Pfam) domain expansions across salmonid genomes. Heatmaps illustrate significant expansions in protein domains (Pfam) across selected salmonid genomes, highlighting enriched domains specifically abundant in (a) *O. formosanus* and (b) *O. m. masou*. Numbers in each cell indicate the total count of the corresponding Pfam domain within each salmonid genome. Colors represent the relative expansion (Z-score) of each domain across species, where positive Z-scores (green to blue shades) indicate higher-than-average domain expansions, and negative Z-scores (pink to red shades) represent lower-than-average copy numbers compared to other salmonids.

**Fig. S12.**
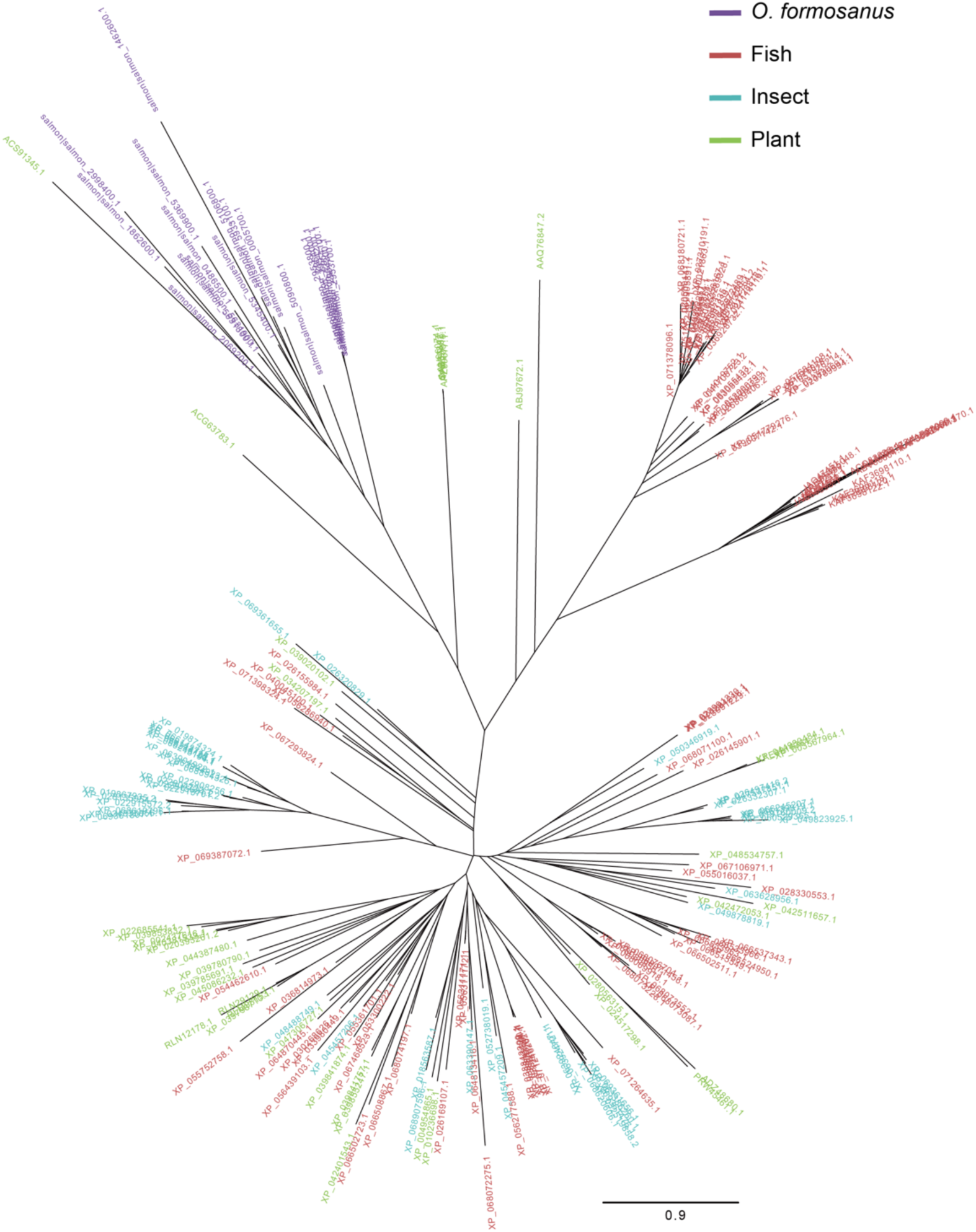
Phylogenetic relationships and divergence of expanded ice-binding lectin domain proteins (LbR_Ice_bind) in *O. formosanus*. Maximum-likelihood phylogeny illustrating the evolutionary divergence of ice-binding lectin proteins (LbR_Ice_bind domain) expanded in *O. formosanus*, compared to characterized ice-structuring protein (ISP) types I, II, and IV (Bar Dolev et al. 2016a) identified in other *Oncorhynchus* species. The distinct cluster formed by *O. formosanus* sequences highlights potential lineage-specific functional specialization associated with adaptation to cold freshwater environments.

**Fig. S13.**
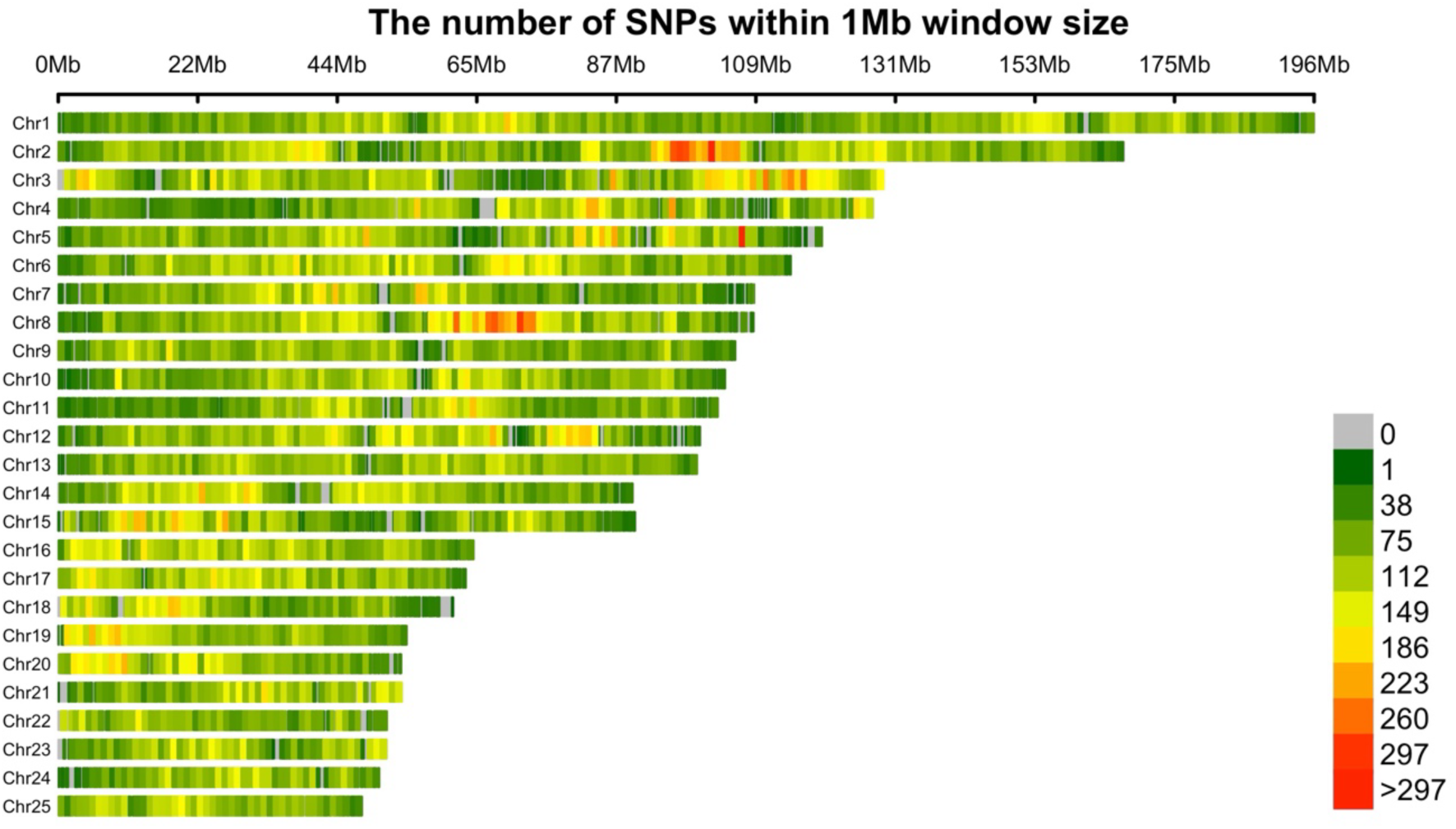
ddRAD-seq mapping results across the genome of *O. formosanus*. SNPs were identified across the newly assembled *O. formosanus* genome using ddRAD-seq data with the window size of 1Mb. The mapping results show even genome-wide coverage, indicating the ddRAD-seq data effectively mapped variation across most regions of the reference genome.

**Fig. S14.**
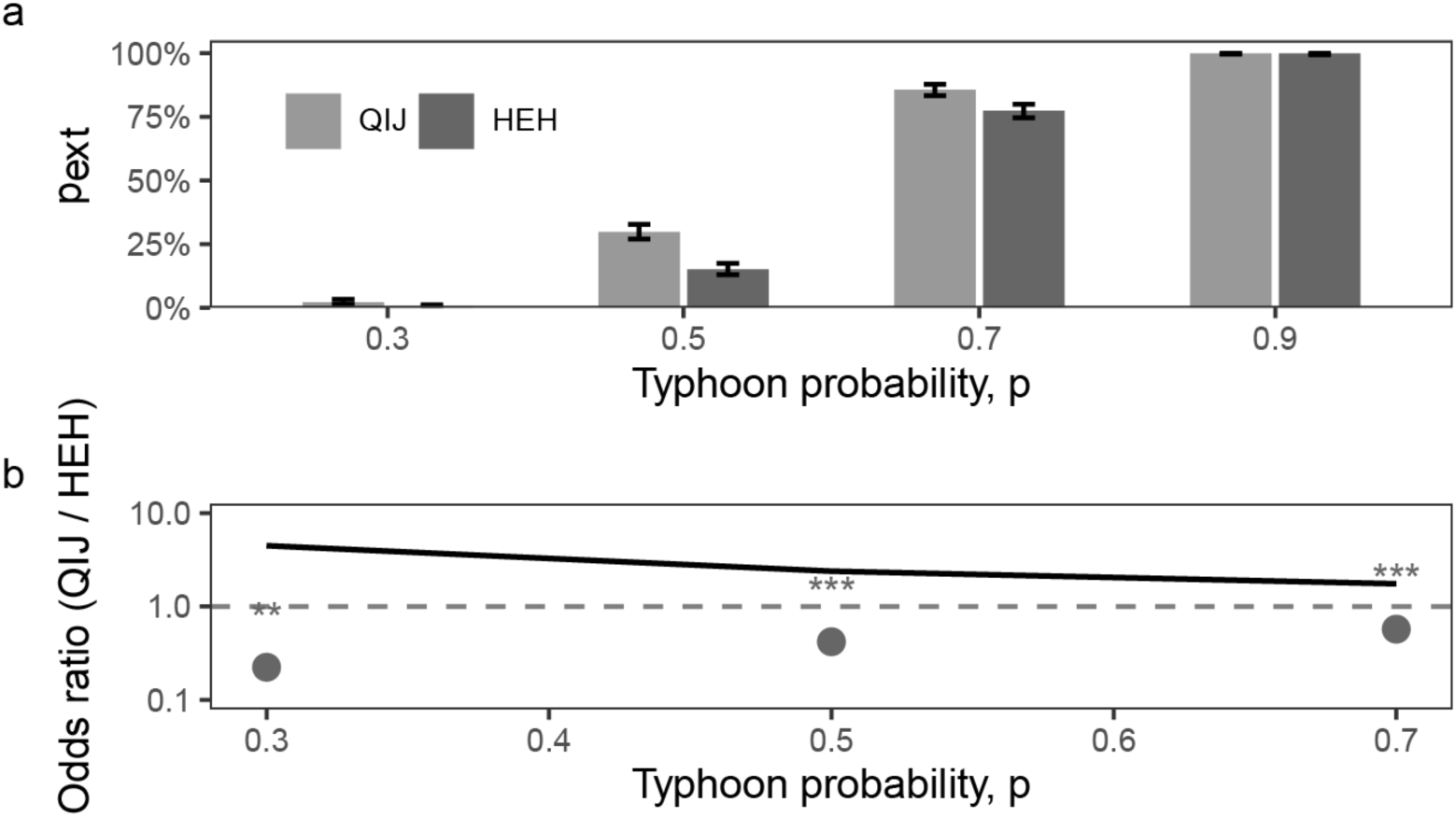
Quasi-extinction probabilities and odds ratios under increasing typhoon disturbance. (a) Quasi-extinction probability of Qijiawan (QIJ, light gray) and Hehuan (HEH, dark gray) across four typhoon frequencies. Bars represent means of 1,000 replicates; whiskers indicate exact 95% binomial confidence intervals. (b) Extinction odds ratios (QIJ / HEH) across typhoon scenarios. Gray points and error bars show Fisher’s exact test results with exact 95% confidence intervals; asterisks indicate significance levels (p < 0.05 *, p < 0.01 **, p < 0.001 ***, ns = not significant). The black line represents logistic GLM estimates of OR across scenarios, showing a monotonic decline with increasing typhoon frequency. The OR at p = 0.9 is omitted due to complete extinction in both populations.

**Fig. S15.**
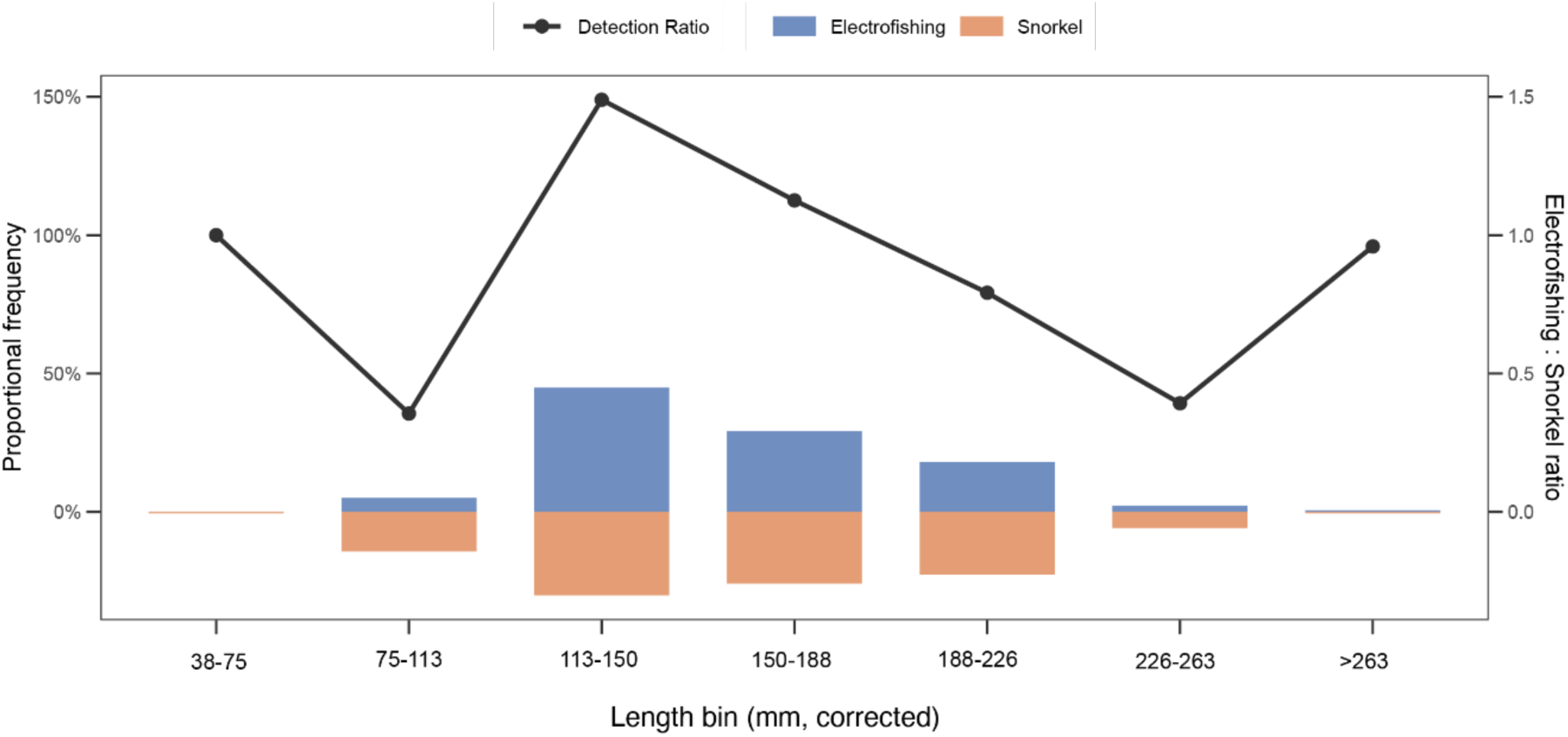
Electrofishing vs. snorkel sampling frequency and detection ratio by length bin. Stacked bars show the proportional frequency of fish in each corrected length bin (mm) as detected by electrofishing (purple) and snorkel surveys (orange) during the same sampling period in 2024 at Hehuan Creek. The green line with markers indicates the detection ratio (electrofishing frequency ÷ snorkel frequency) for each length bin, plotted against the secondary y-axis. Bins with a detection ratio >1 suggest underestimation by snorkel sampling, while ratios <1 suggest potential overestimation.

**Fig. S16.**
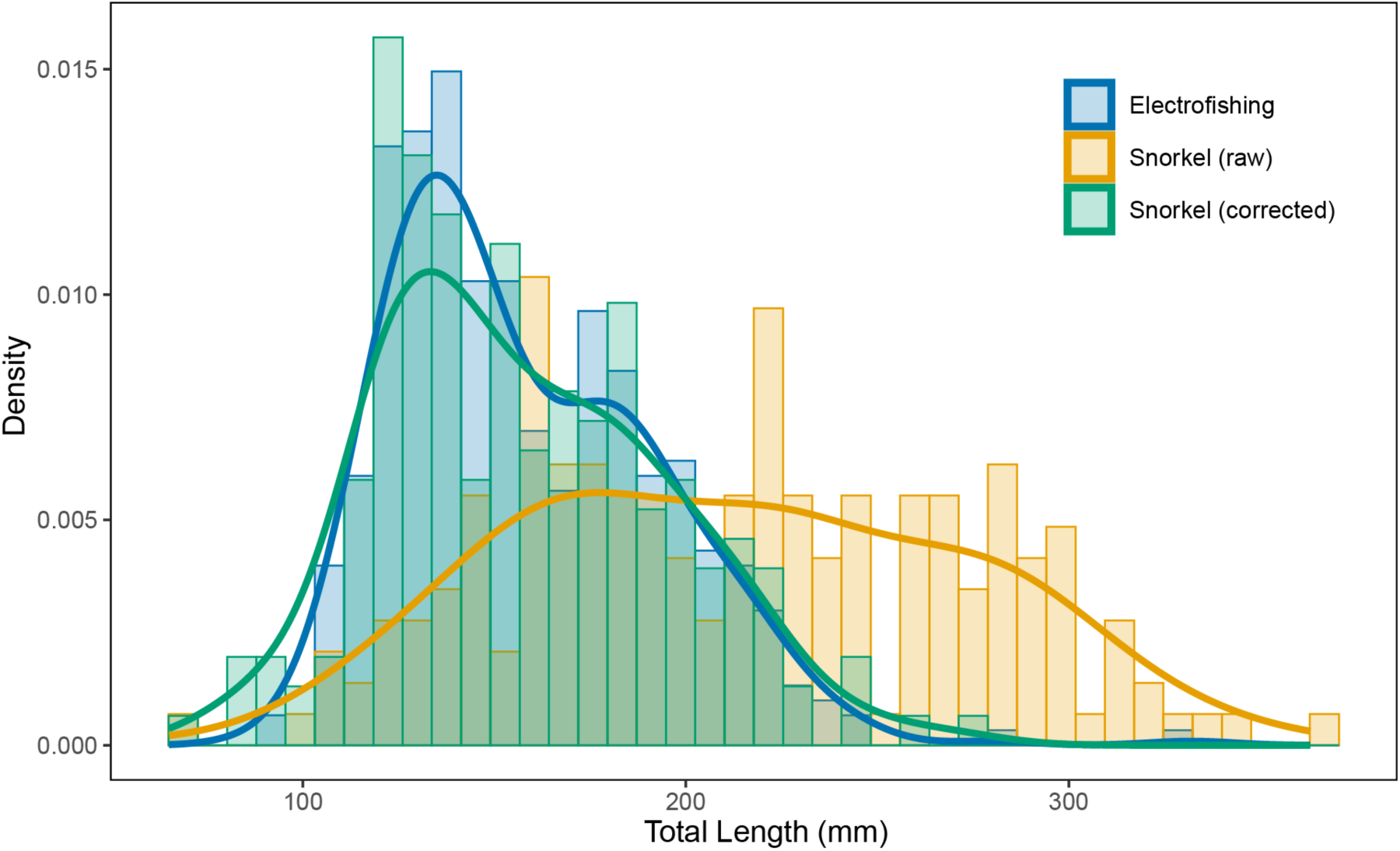
Total length distributions of electrofishing, raw snorkel, and corrected snorkel samples in Hehuan Creek. Histograms show fish total length distributions (mm) obtained via electrofishing (blue), unadjusted snorkel observations (orange), and corrected snorkel estimates (green) during the 2024 sampling period in Hehuan Creek. Corrections for snorkel-derived lengths were applied using refractive index adjustment and detection ratio-based resampling to account for observational biases across length classes.

**Fig. S17.**
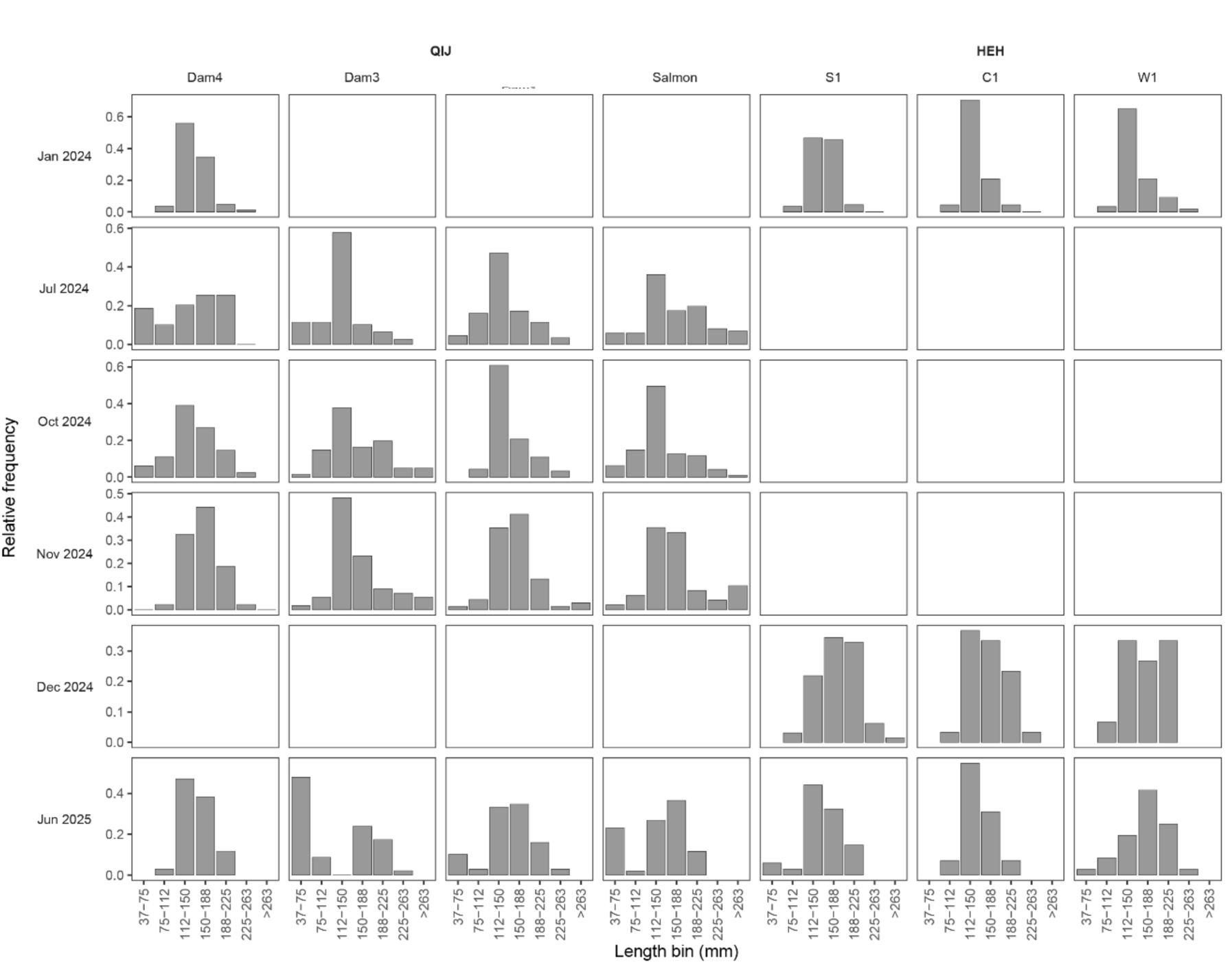
Monthly body-length distributions of Formosan land-locked salmon in two creeks (QIJ and HEH). Relative-frequency histograms (grey bars) show the proportional length composition (mm) of snorkel-derived counts—after detection-ratio and refraction corrections—at seven study sites (Dam4, Dam3, Dam2, Salmon, S1, C1 and W1). Columns are ordered left-to-right by site within each tributary; the tributary names (QIJ, HEH) appear once above their respective site block. Rows correspond to individual monthly surveys (Jan, Jul, Oct, Nov, Dec 2024 and Jun 2025). Bars within each panel sum to 1, enabling direct comparison of length-class structure between site-months regardless of sample size. Blank panels indicate months or sites with no observations.

**Fig. S18.**
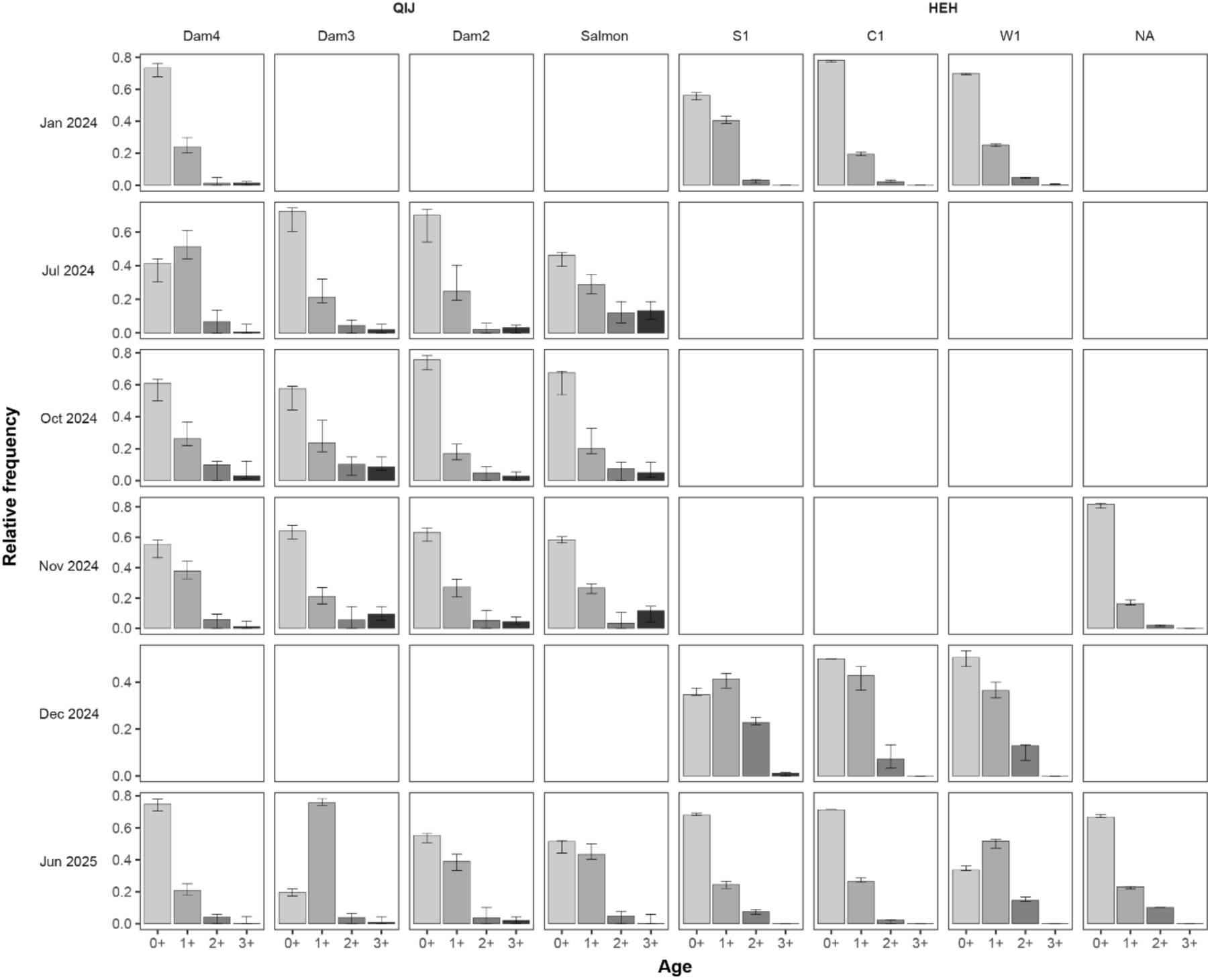
Monthly age-class composition of Formosan land-locked salmon in the QIJ and HEH tributaries of Hehuan Creek. Stacked bars (shaded grey from light to dark for age-classes 0, 1, 2 and 3, respectively) show the posterior mean relative frequency of each age-class at seven study sites (Dam4, Dam3, Dam2, Salmon, S1, C1 and W1). Columns are ordered left-to-right by site within each tributary block (stream labels appear once above the block), while rows correspond to individual monthly surveys (January, July, October, November and December 2024, and June 2025). T-shaped lines denote the 95 % Bayesian credible interval around each bar. Within every panel the four bars sum to 1, allowing direct comparison of age structure across site-month combinations irrespective of sample size. Blank panels indicate months or sites with no observations.

